# Excessive ENaC-mediated sodium influx drives NLRP3 inflammasome-dependent autoinflammation in cystic fibrosis

**DOI:** 10.1101/458208

**Authors:** Thomas Scambler, Heledd H. Jarosz-Griffiths, Samuel Lara-Reyna, Shelly Pathak, Chi Wong, Jonathan Holbrook, Fabio Martinon, Sinisa Savic, Daniel Peckham, Michael F. McDermott

## Abstract

Cystic Fibrosis (CF) is caused by mutations in the cystic fibrosis transmembrane conductance regulator (CFTR) gene and results in defective CFTR-mediated chloride transport, dysregulation of epithelial sodium channels (ENaC) and exaggerated innate immune responses. We tested the hypothesis that upregulation of ENaC drives autoinflammation in this complex monogenic disease.

We show that monocytes from patients with CF exhibit a systemic proinflammatory cytokine signature, with associated anti-inflammatory M2-type macrophage deficiency. Cells harboring CF mutations are hyperresponsive to NLRP3 stimulation, as evidenced by increased IL-18, IL-1β, ASC-specks levels in serum and caspase-1 activity in monocytes, and by increased IL-18 production and caspase-1 activity in human bronchial epithelial cells (HBECs). In both cell types there is an associated shift to glycolytic metabolism with succinate release, in response to increased energy requirements. Inhibition of amiloride-sensitive sodium channels partially reverses the NLRP3-dependent inflammation and metabolic shift in these cells. Overexpression of β-ENaC, in the absence of CFTR dysfunction, increases NLRP3-dependent inflammation, indicating a CFTR-independent ENaC axis in CF pathophysiology. Sodium channel modulation provides an important therapeutic strategy to combat lung inflammation in CF.

## Introduction

Cystic fibrosis (CF) is the most common life-threatening autosomal recessive disease to affect Caucasian populations. Mutations in the cystic fibrosis transmembrane conductance regulator (CFTR) result in reduced expression, function and intracellular localisation of the CFTR protein (Elborn, 2016, Rowe, Miller et al., 2005).

Clinical manifestations of this debilitating condition include repeated pulmonary infections, innate immune-driven episodes of inflammation (Bals, Weiner et al., 1999, Filkins & O’Toole, 2015, Montgomery, Mall et al., 2017, Whitsett & Alenghat, 2015), pancreatic malabsorption (Elborn, 2016), CF-related diabetes (Ntimbane, Comte et al., 2009), malnutrition (Turck, Braegger et al., 2016) and a unique type of inflammatory arthritis (Dixey, Redington et al., 1988). The CFTR protein is widely expressed in a variety of cells and tissues where it acts as an anion channel, by conducting chloride (Cl^−^) and bicarbonate (HCO3-) ions, and also by regulating a range of epithelial transport proteins, including the epithelial sodium channel (ENaC) (Berdiev, Qadri et al., 2009, König, Schreiber et al., 2002, König, Schreiber et al., 2001, Konstas, Koch et al., 2003, Kunzelmann, 2003).

In healthy lungs, CFTR inhibits ENaC and helps maintain normal volume and composition of airway surface liquid (ASL). The absence or reduction in functional CFTR leads to defective CFTR mediated anion transport and upregulation of ENaC. These changes in normal homeostasis result in reduced ASL volume, abnormally thick viscous mucus and defective mucociliary clearance (Althaus, 2013).

While there is no literature to support a direct link between ENaC and inflammation in CF, there is indirect evidence suggesting that aberrant sodium transport influences the disease process. Overexpression of β-ENaC in mice results in a CF-like lung disease, with ASL dehydration, inflammation and mucous obstruction of the bronchial airways (Mall, Grubb et al., 2004a, Zhou, Duerr et al., 2011, Zhou, Treis et al., 2008). In human patients, genetic variants in the *β-* and *γ-ENaC* chains, leading to functional abnormalities in ENaC, have also been associated with bronchiectasis and CF-like symptoms (Fajac, Viel et al., 2008). By contrast, rare mutations, which cause hypomorphic ENaC activity, can slow disease progression in patients homozygous for a deletion of phenylalanine at position 508 in the CFTR (the ΔF508 mutation), which results in failure of the protein to fold into its normal functional conformation (Donaldson & Boucher, 2007, Mall et al., 2004a, Zhou et al., 2011). These studies highlight the essential role of ENaC plays in regulating normal airway homeostasis and show that inhibition of ENaC may modify disease progression either by altering ASL, or modifying other processes, such as inflammation.

The NLRP3 inflammasome has been described as an important inflammatory pathway in CF but there is little indication as to the underlying mechanism(s) and whether this reflects innate autoinflammation or a response to chronic bacterial infections, due to pathogens such as *Pesudomonas aeruginosa* and *Burhkholderia cepacia complex* (Fritzsching, Zhou-Suckow et al., 2015, Iannitti, Napolioni et al., 2016, Montgomery et al., 2017, Rimessi, Bezzerri et al., 2015). NLRP3 and other inflammasomes are activated by various pattern associated- and danger associated-molecular patterns (PAMPs and DAMPs respectively), with the non-canonical inflammasome utilising caspase-11 in mice (Yi, 2017). All of the known inflammasomes, including the NLRP3 inflammasome, function as a signaling platform for caspase-1-driven activation of IL-1-type cytokines (IL-1β and IL-18). Some of IL-18’s main functions include the induction of IFN-γ secretion by natural killer (NK) and T-cells, whereas IL-1β has a role in inducing fever and immune cell proliferation, differentiation and apoptosis. Sustained activation of an inflammasome may induce a caspase-1-dependent form of inflammatory cell death known as pyroptosis (Cookson & Brennan, 2001). Several *in vitro* studies point to K^+^ efflux being an important trigger of NLRP3 activation (Mariathasan, Weiss et al., 2006, Munoz-Planillo, Kuffa et al., 2013, Yaron, Gangaraju et al., 2015).

The aims of this study were to characterise NLRP3 inflammasome activation in CF and to investigate the role of aberrant sodium flux in driving this process. As recent studies on intracellular metabolism in monocyte/macrophages and dendritic cells (DCs) have provided insights on the functioning of these key cells in innate immunity (O’Neill & Pearce, 2016), we also sought to determine the metabolic state of monocyte/macrophages and bronchial epithelial cells (BECs), which are primary cells of interest in patients with CF. We show that both CF sera and immune cells have a proinflammatory IL-1-type cytokine (IL-1β and IL-18) signature associated with inflammasome activation. Both primary monocytes and human bronchial epithelial cell (HBEC) lines, with CF-associated mutations, were shown to hyper-respond to NLRP3 inflammasome stimulation, compared with controls. Increased proinflammatory IL-18 was particularly significant, with comparable levels to that of cells from patients with systemic autoinflammatory disease (SAID). This hyper-responsiveness was associated with increased intracellular Na^+^ influx and K^+^ efflux, with associated increased glycolytic metabolism and succinate release, which could be abrogated by the addition of an NLRP3 inhibitor and inhibition of overactive amiloride-sensitive sodium channels, with amiloride and S18 peptide. Finally, overexpression of β-ENaC in control HBECs, without CFTR mutations, increased IL-18 levels thereby supporting the notion that excessive ENaC-mediated sodium influx drives NLRP3 inflammasome activation in patients with CF.

## Results

### Elevated inflammasome-related cytokine signature in CF sera

To understand the extent of systemic inflammation in CF we measured serum levels of IL-18, IL-1β, IL-1Ra, TNF and IL-6 in patients with CF and compared these to levels in patients with diagnoses of systemic autoinflammatory diseases SAID (genetically and clinically defined), non-CF bronchiectasis (NCFB) and healthy controls (HC) (Fig. 1 A-F). IL-18 and IL-1β are potent inflammatory cytokines that drive systemic inflammation and myeloid cell recruitment to inflamed tissues (Dinarello Charles, 2017, Gabay, Lamacchia et al., 2010, Martin, 2016). In diseases such as SAID, these cytokines drive sterile inflammation, which responds well to therapy with recombinant IL-1Ra. Interestingly, IL-18, IL-1β and IL-1Ra were all significantly elevated in sera from patients with CF, with levels comparable to those found in individuals with SAIDs (Fig. 1 A-C). It is important to note that all the SAID patients in this study were all on active recombinant IL-1Ra (anakinra®) therapy, which will reduce serum IL-1β levels. However, levels of IL-18 and endogenous IL-1Ra are raised in patients with SAID, and are comparable to the proinflammatory, IL-1-type cytokine signature found in CF serum. Notably, other proinflammatory cytokines, associated with innate immune activation and immunological pathologies, including TNF and IL-6 (Fig. 1 E-F), were not elevated to the same extent as key members of the IL-1 cytokine family (Fig. 1 A-C). These data suggest that IL-1 cytokines play a significant role in human CF disease pathology, comparable to SAIDs, but not to any significant degree in NCFB.

**Figure 1:**
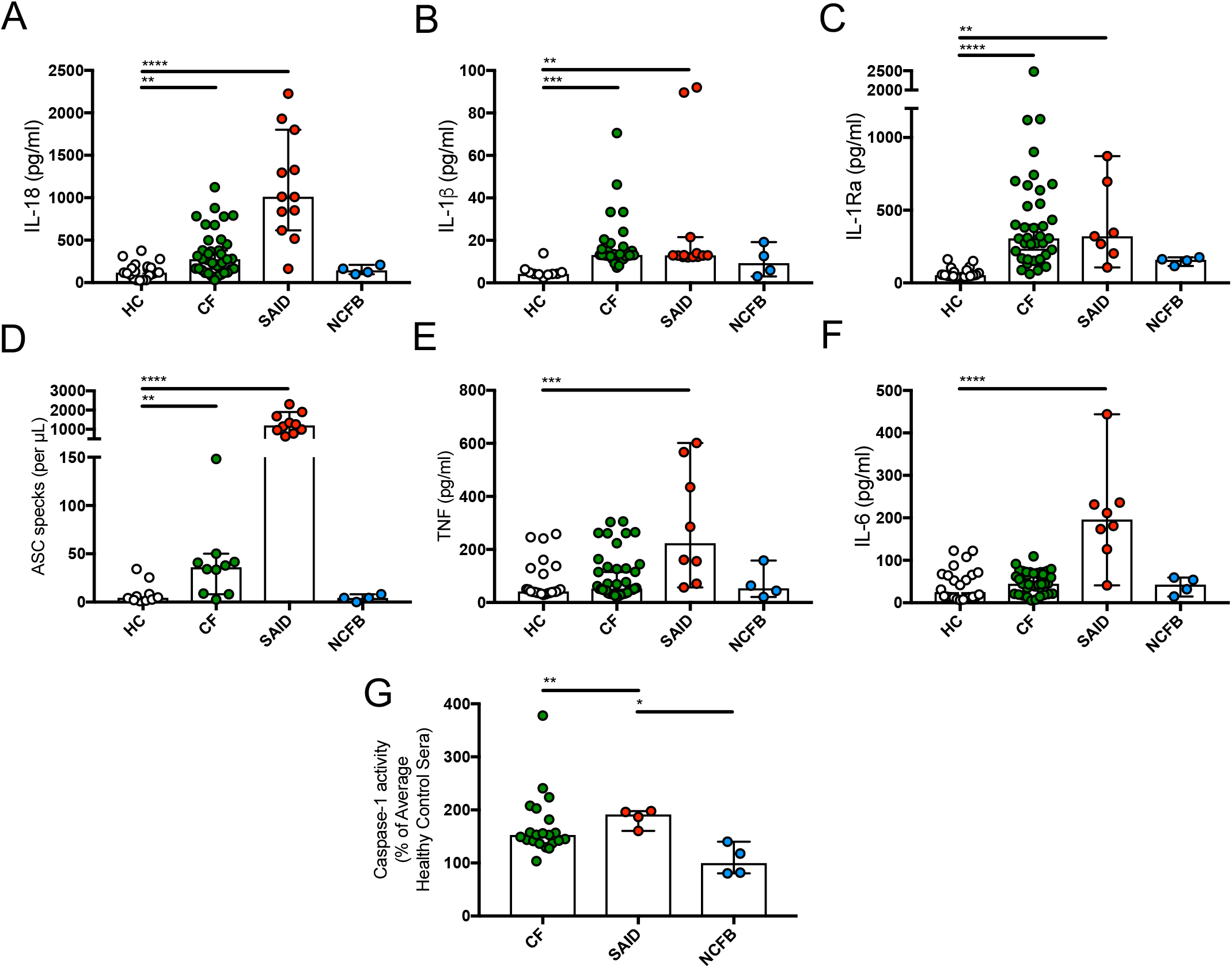
Inflammatory serum cytokine signature in CF. **A-F** ELISA assays were used to detect IL-18 (HC n=21, CF n=36, SAID n=12, NCFB=4) (A), IL-1β (HC n=10, CF n=31, SAID n=13, NCFB=4) (B), IL-1Ra (HC n=20, CF n=37, SAID n=7, NCFB=4) (C), TNF (E) and IL-6 (HC n=24, CF n=40, SAID n=8, NCFB=4) (F) in patient serum. **D** Flow cytometry was used to detect ASC specks in sera of patients with CF, SAID, NCFB and HCs (HC n=10, CF n=10, SAID n=10, NCFB=4). **G** A colorimetric assay to detect caspase-1 activity in sera of patients with CF, SAID and NCFB as a percentage of the average caspase-1 activity from HC (n=10) with a mean absorbance of 0.20845 nm (HC n=10, CF n=21, SAID n=4, NCFB=4). Data Information: The Mann-Whitney non-parametric test was performed (p values * = 0.05, **= 0.01, ***= 0.001 and ****= 0.0001).

Next, we sought to confirm that the raised proinflammatory IL-1-type cytokines in serum are associated with inflammasome activation. We used a flow cytometry-based assay to detect the presence of extracellular apoptosis-associated speck-like protein containing a caspase recruitment domain (ASC) protein aggregates (specks), in sera from HC and patients with CF, SAID and NCFB (Fig. 1 D) (Fernandes-Alnemri & Alnemri, 2008, Franklin, Bossaller et al., 2014). ASC is an integral component of many inflammasomes and when oligomerised, forms large signaling platforms for caspase-1 recruitment. ASC specks are characteristically released from within the cell following inflammasome activation, thereby provoking pyroptotic cell death (He, Wan et al., 2015, Kayagaki, Stowe et al., 2015, Shi, Zhao et al., 2015). ASC specks in sera were found to be significantly elevated in the sera of both CF patients and SAID patients compared to HC (Fig. 1 D). The presence of ASC specks in the serum reflects the presence of NLRP3 inflammasome-mediated inflammation and pyroptosis (Baroja-Mazo, Martin-Sanchez et al., 2014, Fernandes-Alnemri & Alnemri, 2008, Franklin et al., 2014), but ASC filament formation can also serve as an amplification mechanism for a range of different inflammasomes, such as NLRC4 (Dick, Sborgi et al., 2016) and AIM-2 (Matyszewski, Morrone et al., 2018).

To further confirm the presence of inflammasome activation in the peripheral blood of patients with CF, the activity of caspase-1, the rate-limiting factor in the activation of all inflammasomes (Mariathasan et al., 2006, Verhoef, Kertesy et al., 2005), was also measured in serum. Both CF and SAID sera had significantly elevated caspase-1 activity compared to that of NCFB, which was equivalent to HC (Fig. 1 G). Collectively, these data provide robust evidence for inflammasome activation (elevated IL-1 cytokines and ASC specks with increased caspase-1 activity) in patients with CF-associated mutations, with levels comparable to patients diagnosed with SAID.

### Hyper-responsive NLRP3 inflammasome activation in cells with CF-associated mutations

On the basis that proinflammatory IL-1-type cytokines are a hallmark of human CF, we next sought to identify the cellular and molecular sources of IL-18 and IL-1β in the sera. As both IL-18 and IL-1β are typically cleaved and activated by inflammasome-driven caspase-1, we initially explored NLRP3 inflammasome activation in monocytes (the main producers of IL-18 and IL-1β) and HBEC ((BEAS-2B wild-type (WT) control, IB3-1 (ΔF508/W1282X), CuFi-1 (ΔF508/ΔF508) and CuFi-4 (ΔF508/G551D)) lines. To do this, we stimulated primary monocytes with LPS, a broad inflammasome priming signal used to promote pro-IL-18/IL-1β expression, followed by ATP, a specific NLRP3 inflammasome activating signal that assembles NLRP3 with ASC, pyrin and caspase-1 by mediating K^+^ efflux (Fig. 2 A-H). K^+^ efflux is known to act as a potent NLRP3 inflammasome activating signal, common to many NLRP3 inflammasome activators, and can induce NLRP3 inflammasome activation without a priming signal in some circumstances (Munoz-Planillo et al., 2013). Consistent with the serum data, not only did we observe hyper-responsive IL-18 and IL-1β secretion in CF monocytes relative to HC, but when these cells were pretreated with small molecule inhibitor of NLRP3 (MCC950), these secretions were significantly abrogated to the point of being almost undetectable across all patient groups (HC, CF, SAID and NCFB) (Coll, Robertson et al., 2015) (Fig. 2 A-B). Secretion of IL-18 and IL-1β was also significantly decreased when TLR4 and caspase-1 were inhibited by pretreatment with small molecule inhibitors, OxPAC and AC-YVADD-cmv, respectively (Fig. 2 A-B). NLRP3 inflammasome activation in HBECs, measured using IL-18, was also found to be upregulated in the CF-associated mutant cell lines (IB3-1 and CuFi-1) relative to BEAS-2B control, which was reduced with NLRP3, caspase-1 and TLR4 inhibitors (Fig. 2 D). It is important to note that although IL-18 cytokines are released from HBECs in response to inflammasome activation, consistent with monocyte secretion, IL-1β cytokine release by HBECs was not detectable by ELISA. Interestingly, pro-IL-1β gene expression is increased in the CF-associated mutant cells, CuFi-1 and CuFi-4 relative to BEAS-2B control cells at baseline, but LPS stimulation does not increase pro-IL-1? gene transcript levels, as might be expected (Supplementary Fig 1 A). This discrepancy is consistent with previous reports in which IL-18 alone is enough to abrogate NLRP3 driven hyperactivity and also highlights the varying specificities of different cell types for production of IL-1 cytokine family members. Caspase-1 activation was also elevated in CF and SAID monocytes (Fig. 2 C) and compared to both HC and NCFB monocytes and in IB3-1 cells (Fig. 2 E) relative to BEAS-2B control (Fig. 2 E), post LPS and ATP stimulation *in vitro*. Notably, caspase-1 activity in all monocyte and HBEC cohorts was depleted by MCC950 pretreatment (Fig. 2 C and E). In contrast to the serum data, TNF was also elevated in CF monocytes, following LPS and ATP stimulation, but was not modulated by MCC950 (Supplementary Fig. 2 A). To rule out the involvement of any other caspase-1 driven inflammasome, we also stimulated monocytes and HBEC lines with established activators of three other inflammasomes, namely NLRC4, pyrin and AIM-2. There were no statistically significant differences between cells with or without CF-associated mutations for activation of these particular inflammasomes (Supplementary Fig. 1 B-D). In support of elevated IL-1 serum cytokine signature in CF, we have shown that CF monocytes and CF-associated mutant epithelial cells respond excessively to NLRP3 inflammasome stimulation, with IL-18 being a key player across both monocyte and epithelial cells.

**Figure 2:**
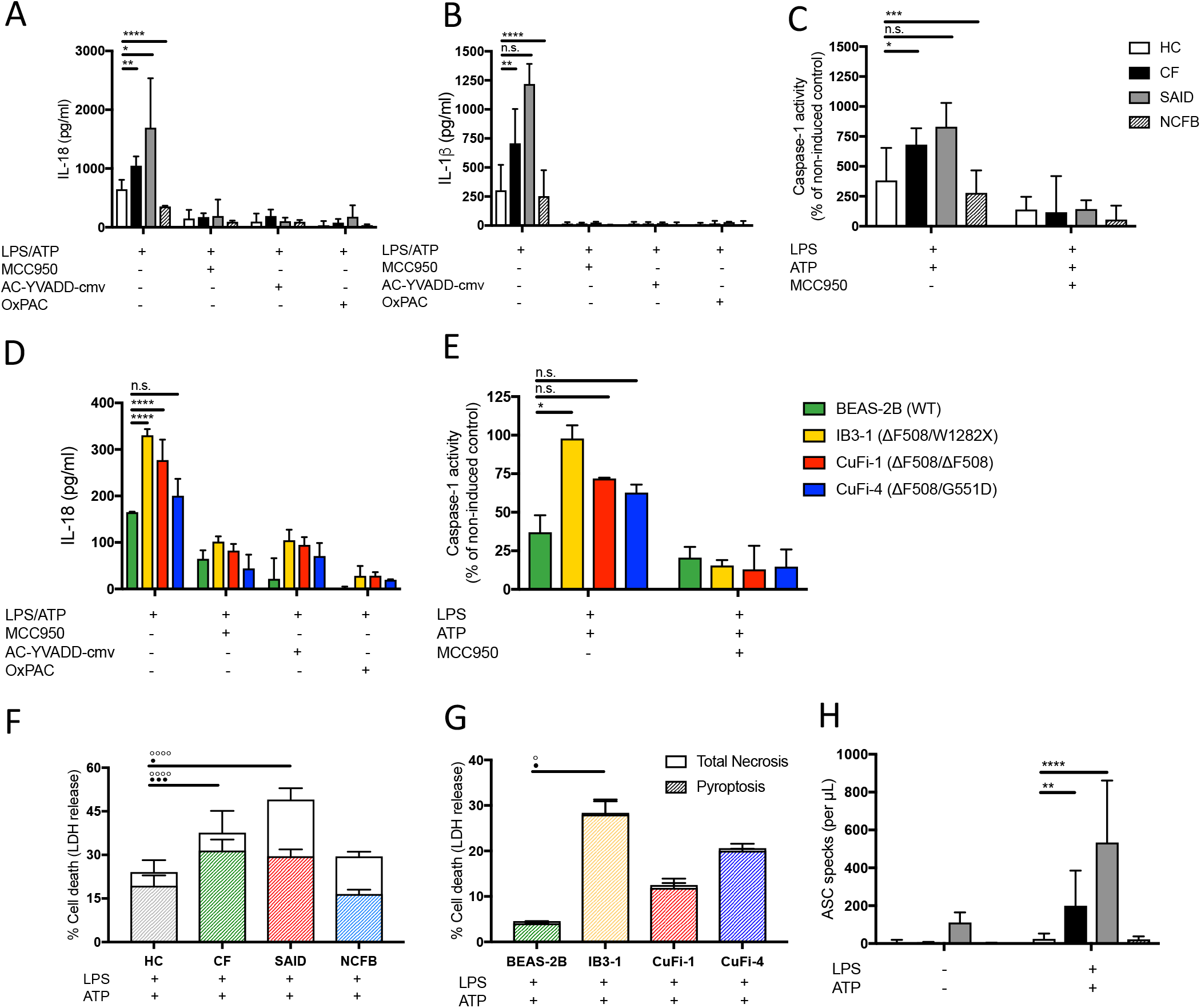
Hyper-inflammation in CF is NLRP3 dependent. **A-E** Primary monocytes from HC, CF, SAID and NCFB (HC n=10, CF n=10, SAID n=4, NCFB=4) (A-C) or HBEC lines (n=3) (D, E) were pre-incubated with MCC950 (15mM), OxPAC (30mg/mL) and Ac-YVADD-cmv (2mg/mL) for 1-hour before a stimulation with LPS (10ng/mL, 4 hours), and ATP (5mM) for final 30 minutes. ELISA assays were used to detect IL-18 (A,D) and IL-1b (B) cytokine secretion in supernatants and colometric assay used to detect caspase-1 activity in protein lysates (C, E). **F-G** Necrosis and pyroptosis are represented as superimposed bar charts (G, H). Total necrosis was measured using LDH release assay. For pyroptotic cell death, each sample/condition was repeated in parallel with a caspase-1 inhibitor (AC-YVADD-cmv (2mg/mL, 1 hour)) pre-treatment. The total necrosis level was then taken away from the capase-1 inhibited sample, or ‘caspase-1 independent’ necrosis, with the remaining LDH level termed ‘caspase-1 dependent necrosis’ or pyroptosis. Cells were then stimulated with LPS (10ng/mL, 4 hours), and ATP (5mM) for final 30 minutes. The assay was performed with primary monocytes from HC, CF, SAID and NCFB (HC n=10, CF n=10, SAID n=4, NCFB=4 and (H) HBEC lines (n=3). (⚬) Significance for Total Necrosis (•) Significance for Pyroptosis. **H** Flow cytometry was used to detect ASC specks in supernatant of primary monocytes from HC, CF, SAID and NCFB (HC n=6, CF n=6, SAID n=6, NCFB=4). Data Information: A 2-way ANOVA statistical test was performed (n.s. = not-significant; p values * = 0.05, **= 0.01, ***= 0.001 and ****= 0.0001). All inhibitor treatments in panels A-E were found to significantly reduce cytokine secretion and caspase-1 activity to **p= 0.01 or less. Significance values not displayed on the graph.

We next examined necrotic cell death in HC, CF, SAID and NCFB monocyte cohorts and HBEC lines (Fig. 2 F-G). Pyroptosis was distinguished from total necrosis by pretreating the cells with a caspase-1 inhibitor and using lactose dehydrogenase (LDH) as a measure of necrosis. Elevated pyroptosis was present after LPS and ATP stimulation in monocytes from patients with CF, and those diagnosed with SAID, as well as the HBEC line IB3-1 (Fig. 2 F-G). Notably, caspase-1-independent necrosis was also elevated in CF and SAID monocytes (Fig. 2 F). To understand the relationship between elevated pyroptosis and downstream inflammation, ASC specks were measured in the cell medium, post-stimulation of monocytes for NLRP3 inflammasome activation with LPS and ATP (Fig. 2 H). ASC specks were elevated in stimulated monocytes from patients with CF-associated mutations and also in those diagnosed with SAID.

An immunological consequence of elevated IL-1β and IL-18 secretion by monocytes is that other immune cells are activated. IL-1β induces proliferation, differentiation and apoptosis (Church, Cook et al., 2008, Gabay et al., 2010, Iannitti et al., 2016, Martin, 2016, Miller, Pietras et al., 2007, Palomo, Dietrich et al., 2015) and IL-18 acts on NK and T-cells to express and secrete IFN-γ (Chaix, Tessmer et al., 2008, Dinarello, 1999, Kim, Chae et al., 2015, Marshall, Aste-Amézaga et al., 1999, van de Veerdonk, Netea et al., 2011). We therefore investigated whether the elevated IL-18 secretion by CF monocytes induced elevated IFN-γ secretion in the peripheral blood mononuclear cell (PBMC) population. We isolated PBMCs and stimulated with LPS and ATP to activate the NLRP3 inflammasome within the monocytes. IFN-γ gene expression and secretion were significantly increased, post-NLRP3 inflammasome activation, in PBMCs from patients with CF compared to HC (Supplementary Fig. 1 E-F). These findings further indicate that NLRP3 inflammasome activation disseminates local and systemic inflammation, via pyroptosis and activation of IFN-γ producing cells, in patients with CF.

### Classical monocyte and M2-type macrophage populations are reduced in CF

To determine if the elevated serum and monocyte IL-1β and IL-18 levels observed in CF (Fig. 1) correlate with a chronic, inflammatory cellular response, we next studied the capacity of monocytes to differentiate into M1-type and M2-type macrophages. M1-type macrophages are considered to have a proinflammatory role in peripheral tissues, and are characterized by an inherent ability to produce proinflammatory IL-6 and TNF cytokines, whilst M2-type macrophages are archetypically associated with production of the anti-inflammatory cytokine, IL-10 (Italiani & Boraschi, 2014, Meyer, Huaux et al., 2009, Mills, Kincaid et al., 2000, Van den Bossche, Baardman et al., 2015). Peripheral monocytes were differentiated and activated into either M1-type or M2-type macrophages (Italiani & Boraschi, 2014, Mia, Warnecke et al., 2014) and characterised by measuring macrophage cell surface markers and intracellular cytokines using flow cytometry and ELISA (Fig. 3 A-F) (Mills et al., 2000, Tarique, Sly et al., 2017). Both M1-type and M2-type macrophages have distinct surface markers (Supplementary Fig. 3 A) and morphology (Supplementary Fig. 4 A). Following *ex vivo* monocyte differentiation, we observed a statistically significant deficiency in the ability of primary monocytes to differentiate into M2-type macrophages in patients with CF, in comparison with HCs (Fig. 3 B), without any statistical difference being observed in the proportion of M1-type macrophages in either CF or HC cohorts (Fig. 3 C). Despite this, a statistically significant increase in IL-6 was detected in samples originating from CF patients (Fig. 3 D), which indicates that M1-type macrophages from patients with CF-associated mutations are hyper-responsive to inflammatory stimuli. As expected, reduced IL-10 secretion was observed in the monocyte populations, cultured for M2-type macrophage differentiation from these patients (Fig. 3 E).

**Figure 3:**
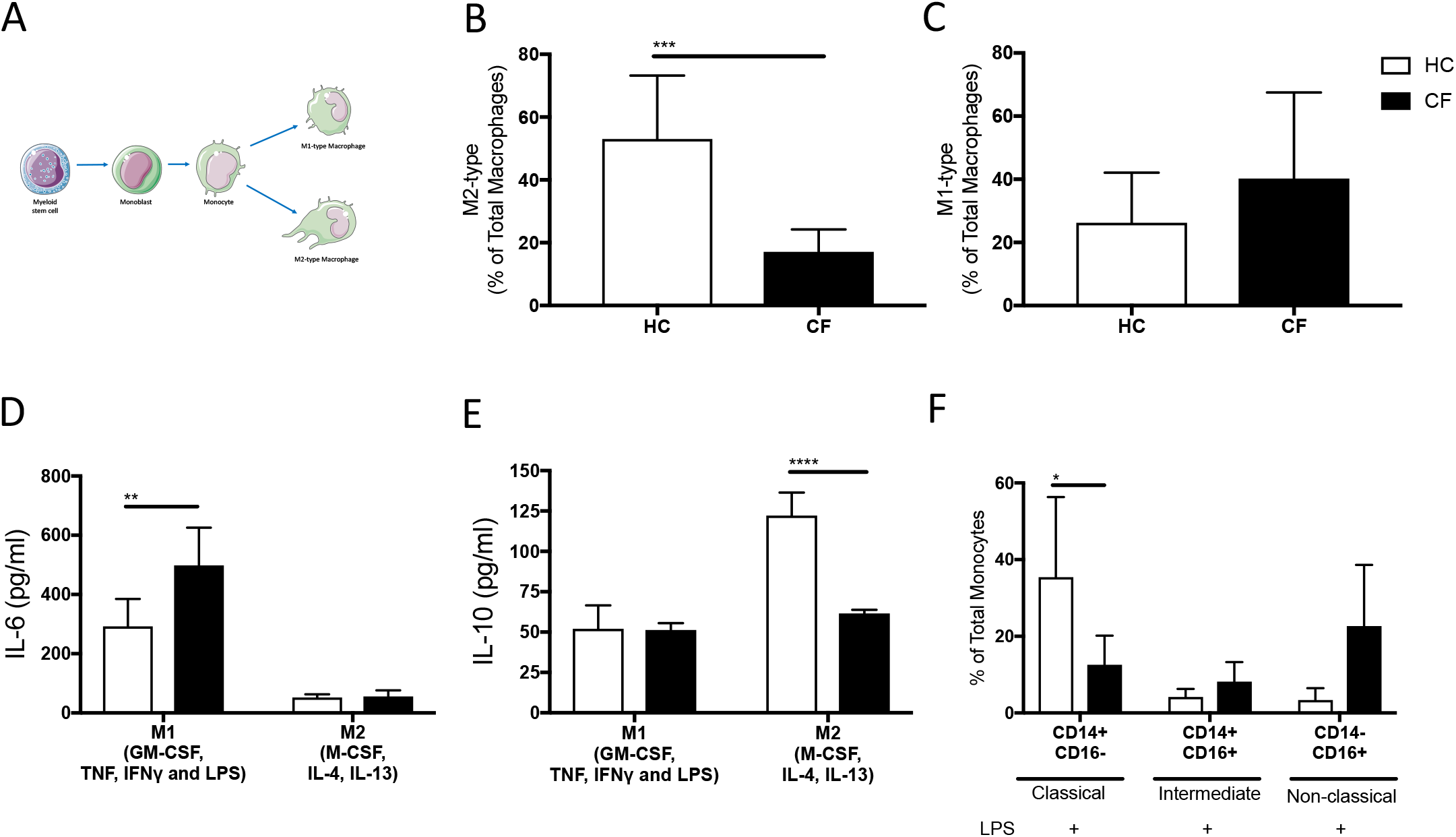
CF monocytes and macrophages have an inflammatory phenotype.] **A** Monocytes were gated on based on forward and side scatter and then based on CD14 and CD16 expression as follows: classical (CD14++CD16−), non-classical (CD14dimCD16++), and intermediate (CD14++CD16+). Cells were stimulated with LPS (10ng/mL, 4 hours). **B** Surface markers for M1-type (CD14+ CD16+ HLA-DR+ CD274+ CD86+) and M2-type (CD14+ CD16+ CD206+) were analyzed using flow cytometry. M1-type macrophage activation was achieved by supplementing growth media with 100ng/mL human IFN-g, 100ng/mL TNF and 50ng/mL LPS. M2-type macrophage activation was achieved by supplementing growth media with 20ng/mL IL-13 and 20ng/mL IL-4. Monocytes from whole blood were differentiated into macrophages and characterised as M1-type (markers-CD14^+^ CD16^+^ HLA-DR^+^ CD274^+^ CD86^+^ TNF^HI^) or M2-type (markers-CD14^+^ CD16^+^ CD206^+^ IL-10^HI^). **C-D** Numbers of polarised macrophages are presented as percentage of total macrophages in samples (n=7). **E-F** ELISA assays were used to detect IL-6 (E) and IL-10 (F) levels in macrophage supernatant (n=7). Data Information: The Mann-Whitney non-parametric test was performed (p values * = 0.05, **= 0.01, ***= 0.001 and ****= 0.0001).

We next examined the phenotype of peripheral monocytes, before their differentiation into M1-type and M2-type macrophages (Fig. 3 A, F), based on their expression of CD14 and CD16 (Supplementary Fig. 3 B). Reduced numbers of expanded, peripheral monocytes of the classical phenotype, known to be phagocytic in function (Patel, Zhang et al., 2017) (Fig. 3 F) was observed. This population of monocytes is not thought to be associated with immunological disease but instead is considered to have a mainly protective effect against bacterial infection (Gordon & Taylor, 2005, Italiani & Boraschi, 2014, Patel et al., 2017, Sprangers, Vries et al., 2016). There was no statistically significant increase in peripheral monocytes with an intermediate or non-classical phenotype associated with chronic inflammatory conditions, such as atherosclerosis and rheumatoid arthritis (Patel et al., 2017, Sprangers et al., 2016) (Fig. 3 F).

Collectively, these data reveal evidence of a deficiency in both classical monocytes and M2-type macrophages; however, no statistically significant increase in proinflammatory monocyte/macrophage populations was observed, and the notable lack of anti-inflammatory macrophages may be responsible for the marked chronicity of the inflammatory response in CF (Bruscia, Zhang et al., 2009, Gordon & Taylor, 2005, Italiani & Boraschi, 2014, Mills et al., 2000).

### Increased metabolism in cells with CF-associated mutations

When immune cells become activated into effector cells (e.g. monocytes to macrophages), there is an attendant adjustment of cellular metabolism, with expression of various transcription factors, cell surface proteins and cytokine secretion, which serve to support the new effector functions of the activated cells (Ganeshan & Chawla, 2014). Cellular metabolism produces energy in the form of ATP through either oxidative phosphorylation (OXPHOS) for either slow, efficient energy production, in the form of ATP, or glycolysis, which supplies a rapid but low-yield of ATP (Buck, Sowell et al., 2017, Ganeshan & Chawla, 2014). Effector cells characteristically utilise glycolysis to fuel acute effector cell functions, so, therefore, cellular metabolism is considered to be a marker of inflammation (Kelly & O’Neill, 2015, Mills, Kelly et al., 2017, O’Neill, 2015).

The extracellular acidification rate (ECAR), which quantifies changes in lactic acid and H^+^ efflux as a byproduct of glycolysis, was used to assess the degree of glycolysis and OXPHOS was used to measure the oxygen consumption rate (OCR), to quantify oxygen depletion in the cell culture medium as cells absorb oxygen for use in the electron transport chain (ETC) and mitochondrial metabolism. We found significantly elevated ECAR in CF monocytes and the CuFi-4 HBEC line after LPS stimulation, to induce an inflammatory phenotype (Fig. 4 A, D). OCR was significantly elevated in IB3-1 and CuFi-1 HBEC lines (Fig. 4 E) but no difference was seen between HC and CF monocytes (Fig. 4 B). Both glycolysis and OXPHOS produce intracellular ATP and this intracellular nucleoside triphosphate was elevated in both monocytes and HBEC lines with CF-associated mutations (Fig. 4 C, F). Together, these data suggest that CF-associated mutations have an intrinsic role in predisposing monocytes and HBECs towards a highly metabolic and inflammatory phenotype.

**Figure 4:**
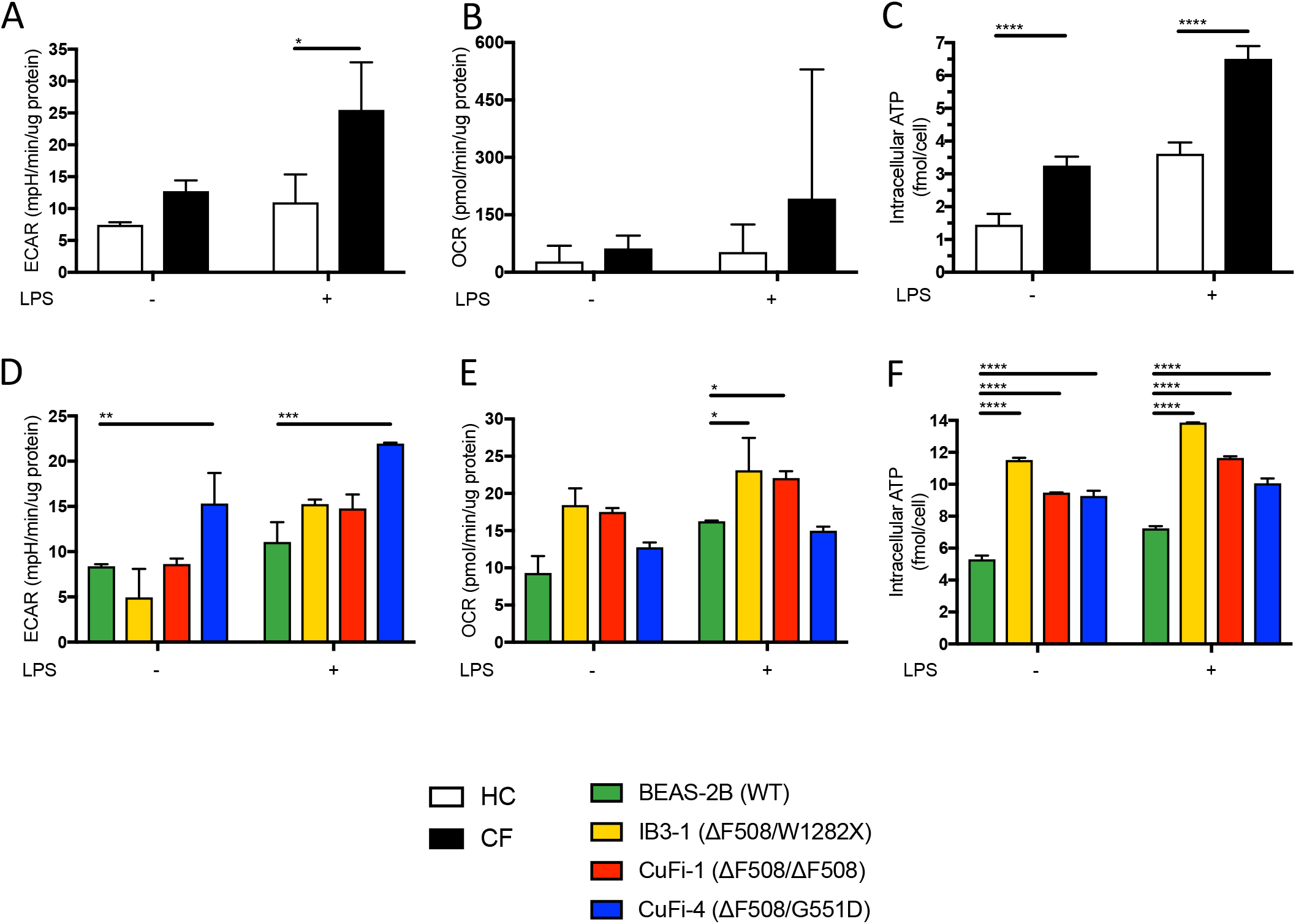
Increased cellular metabolism in cells with CF-associated mutations. **A, B, C, D** Seahorse extracellular flux analyser measured (A and C) the extracellular acidification rate (ECAR) and (B and D) oxygen consumption rate (OCR) in HBECs (n=3), monocytes (n=4) and T-cells (n=4). Oligomycin (10μM), an inhibitor of ATP synthase, and the uncoupling agent, Carbonyl cyanide-4-(trifluoromethoxy) phenylhydrazone (FCCP) (5μM), were used to metabolically stress and depolarise cellular mitochondrial membranes. Cells were pre-treated with amiloride (100mM, 1 hour) before LPS (10ng/mL, 4 hours) stimulation. **E, F** Colorimetric assays were used to detect intracellular ATP in HBECs (n=3) and primary monocytes (n=7). Cell were pre-treated with amiloride (100mM, 1 hour) before an LPS (10ng/mL, 4 hours) stimulation. Data Information: A 2-way ANOVA was performed (p values * = ≤0.05, **= ≤0.01, ***= ≤0.001 and ****= ≤0.0001).

### Increased sodium influx in cells with CF-associated mutations

Next, we explored the inflammatory and metabolic CF phenotype and various mechanisms regulating this process. As K^+^ efflux is known to act as a potent NLRP3 inflammasome activating signal we sought to establish whether K^+^ efflux was also dysfunctional in human CF, thereby leading to excessive NLRP3 activation and signaling. As the presence of overactive amiloride-sensitive Na^+^ channels are characteristic of human and animal models of CF (Mall et al., 2004a, Zhou et al., 2011, Zhou et al., 2008), the resulting increased intracellular Na^+^ cation concentration may be responsible for modulating any increase in K^+^ efflux (Munoz-Planillo et al., 2013). Intracellular Na^+^ influx was significantly higher in CF cells (both monocytes and HBEC), on stimulation with LPS and ATP (Fig. 5A, C), and this was matched by increased efflux of the K^+^ cation (as monitored by a reduction in intracellular K^+^) (Fig. 5 B, D). The magnitude of these fluxes was significantly reduced by the addition of amiloride, a short-acting small-molecule ENaC inhibitor (Fig. 5 A-D). Other small molecule inhibitors (EIPA and S18 peptide) were also used to fully elucidate the specific amiloride-sensitive channels responsible for the differential Na^+^ and K^+^ fluxes in these cells (Fig. 5 A-D). EIPA is able to inhibit broad-spectrum sodium channels, but with lower potency for ENaC. Pretreatment with EIPA reduced Na^+^ influx and K^+^ efflux in CF monocytes and HBEC lines, but not to the same extent as amiloride (Fig. 5 A-D). The SPLUNC1-derived peptide, S18, is a highly stable and specific small molecule inhibitor of ENaC channels. S18 significantly reduced Na^+^ influx and K^+^ efflux in HC and CF monocytes and HBEC lines (Fig. 5 A-D). Ouabain is a small molecule, which inhibits the Na^+^/K^+^ ATP pump. Pretreatment with ouabain increased Na^+^ influx and decreased K^+^ efflux, as expected in both monocytes and HBEC lines and was, therefore, used as a positive control (Fig. 5 A-D).

**Figure 5:**
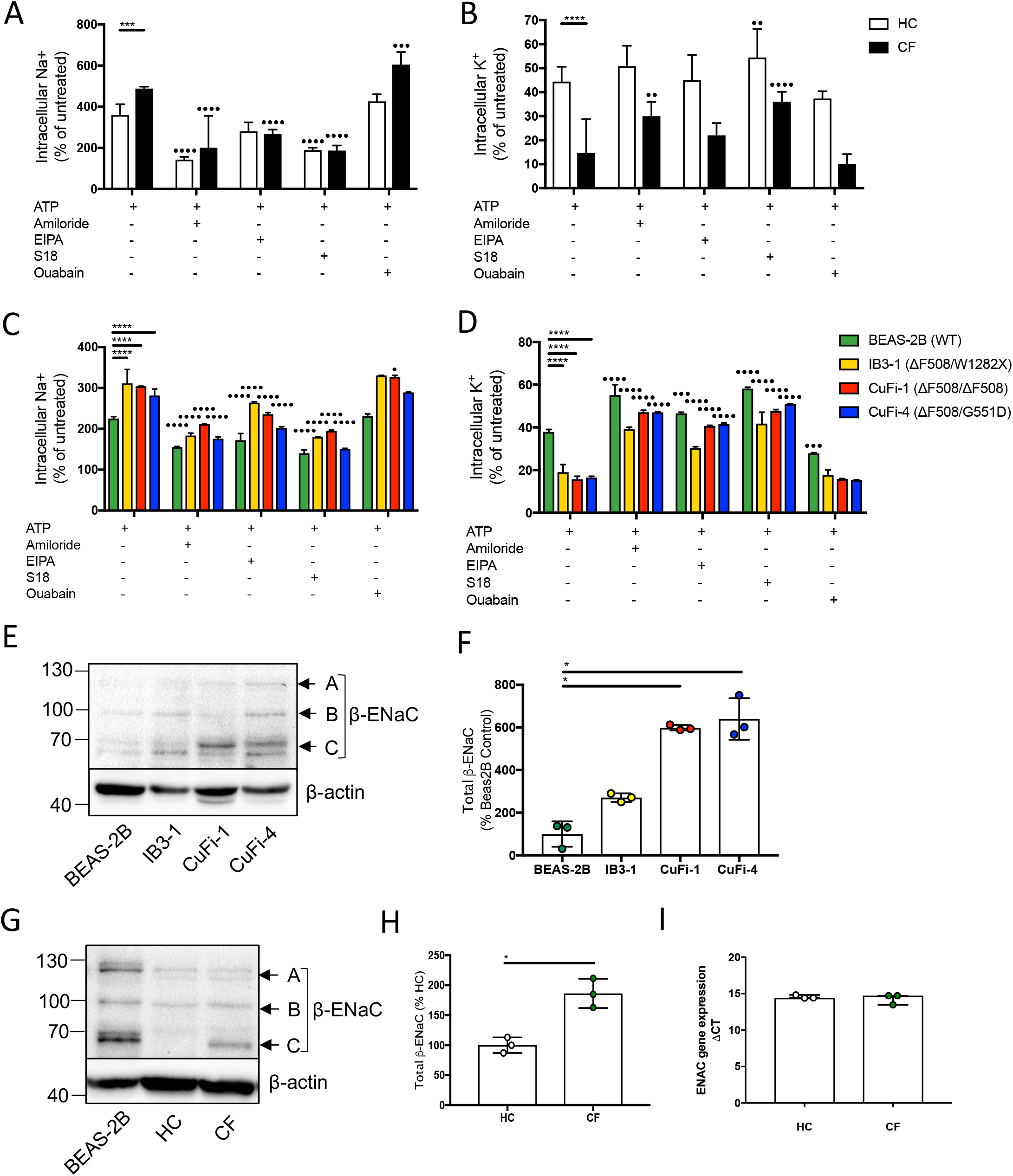
Increased sodium influx in cells with CF-associated mutations. **A-D** Intracellular Na^+^ was detected using an AM ester of potassium indictor PBFI (P-1266) (A, C) and intracellular K^+^ was detected using an AM ester of sodium indictor SBFI (S-1263) (B, D); changes in fluorescence were measured by fluorimeter post-stimulation in monocytes (A, B) from patients with CF (n=7) as well as HBECs (n=3) (C, D). Monocytes and HBECs were pre-treated with the following: amiloride (100mM, 1hour), EIPA (10mM, 1 hour), S18 derived peptide (25mM, 4 hours) and ouabain (100nM, 24 hours) before a stimulation with LPS (10ng/mL, 4 hours) and ATP (5mM) for the final 30 minutes. **E-F** Endogenous β-ENaC protein expression was detected using western blot in BEAS-2B, IB3-1, CuFi-1 and CuFi-4 HBEC lines (E) and densitometry analysis of total β-ENaC (bands A, B, C indicated on blot) was quantified in (F) (n=3). Band A represents complex N-Glycosylation, 110kDa β-ENaC (found when associated as ENaC complex); Band B represents Endo-H sensitive N-Glycosylation, 96kDa β-ENaC; Band C represents immature un-glycosylated, 66kDa β-ENaC. **G-H** Endogenous β-ENaC protein expression was detected using western blot in BEAS-2B HBEC, HC and CF monocytes (G) and densitometry analysis of total β-ENaC (bands A, B, C indicated on blot) was quantified in (H) for CF relative to HC (n=3). **I** Gene expression of β-ENaC in HC vs CF (n=1), represented as DCT. Data Information: A-D A 2-way ANOVA statistical test was performed (p values *= 0.05, **= 0.01, ***= 0.001 and ****= 0.0001); E-H The Mann-Whitney non-parametric test was performed (p values * = 0.05). (*) indicate significance when comparing HC with CF. (•) indicate significance between treatments within the same cell line.

As inhibition of ENaC is able to modulate not only Na^+^ influx but also K^+^ efflux upon ATP stimulation of the P2X7 receptor, we measured β-ENaC protein expression in HBEC lines and found a significant increased expression in CuFi-1 and CuFi-4 cells relative to BEAS-2B control (Fig. 5 E-F). We show, for the first time, that β-ENaC is expressed at the gene expression level in monocytes (Fig. 5 I) and also at the protein level (Fig. 5 G-H) with a significant increase in β-ENaC in CF monocytes relative to HC at the protein level (Fig 5 G-H).

These data demonstrate that β-ENaC is expressed to a greater extent in CF HBECs and CF monocytes; not only is this channel expressed and functional, but is also overactive in monocytes with CF-associated mutations. Interestingly, inhibition of Na^+^ influx with specific small molecules and peptides is also able to also modulate K^+^ efflux, on ATP stimulation of the P2X7 channel. These findings indicate that elevated Na^+^ influx in both monocytes and HBECs with CF-associated mutations may drive NLRP3 inflammasome activation by reducing the required threshold for K^+^ efflux, a potent NLRP3 inflammasome activating signal.

### Inhibition of amiloride-sensitive sodium channels modulates inflammation in CF

Subsequent experiments examined the extent to which dysregulated Na^+^ flux influences cellular metabolism in patients with CF, and if this dysregulation contributes to the observed NLRP3 inflammasome activation. Notably, amiloride alleviated the already augmented cytokine secretion (Fig. 6 A-C) and caspase-1 activity (Fig. 6 J-K) in both primary CF monocytes and HBEC lines, consistent with previously reported murine studies using the β-ENaC Tg-mouse (Fritzsching et al., 2015, Mall et al., 2004a, Mall, Grubb et al., 2004b, Montgomery et al., 2017). EIPA was used to elucidate whether ENaC was central to this amiloride-sensitive channel dependent NLRP3 inflammasome over-activation and we found that EIPA was unable to modulate IL-18, IL-1β or caspase-1 activity in monocytes or HBECs from any cohort (Fig. 6 D-F). Interestingly, the SPLUNC1-derived peptide, S18 potently inhibited cytokine secretion and caspase-1 activity exclusively in cells with CF-associated mutations, as was the case with amiloride (Fig. 6 G-I). These data were replicated by nigericin activation of NLRP3 as a control for ATP, due to ATP’s ability to modulate other ionic channels (Supplementary Fig. 5 A-I). Finally, inhibition of amiloride-sensitive channels did not modulate TNF (Supplementary Fig. 2 C), suggesting a specific ENaC-NLRP3 axis.

**Figure 6:**
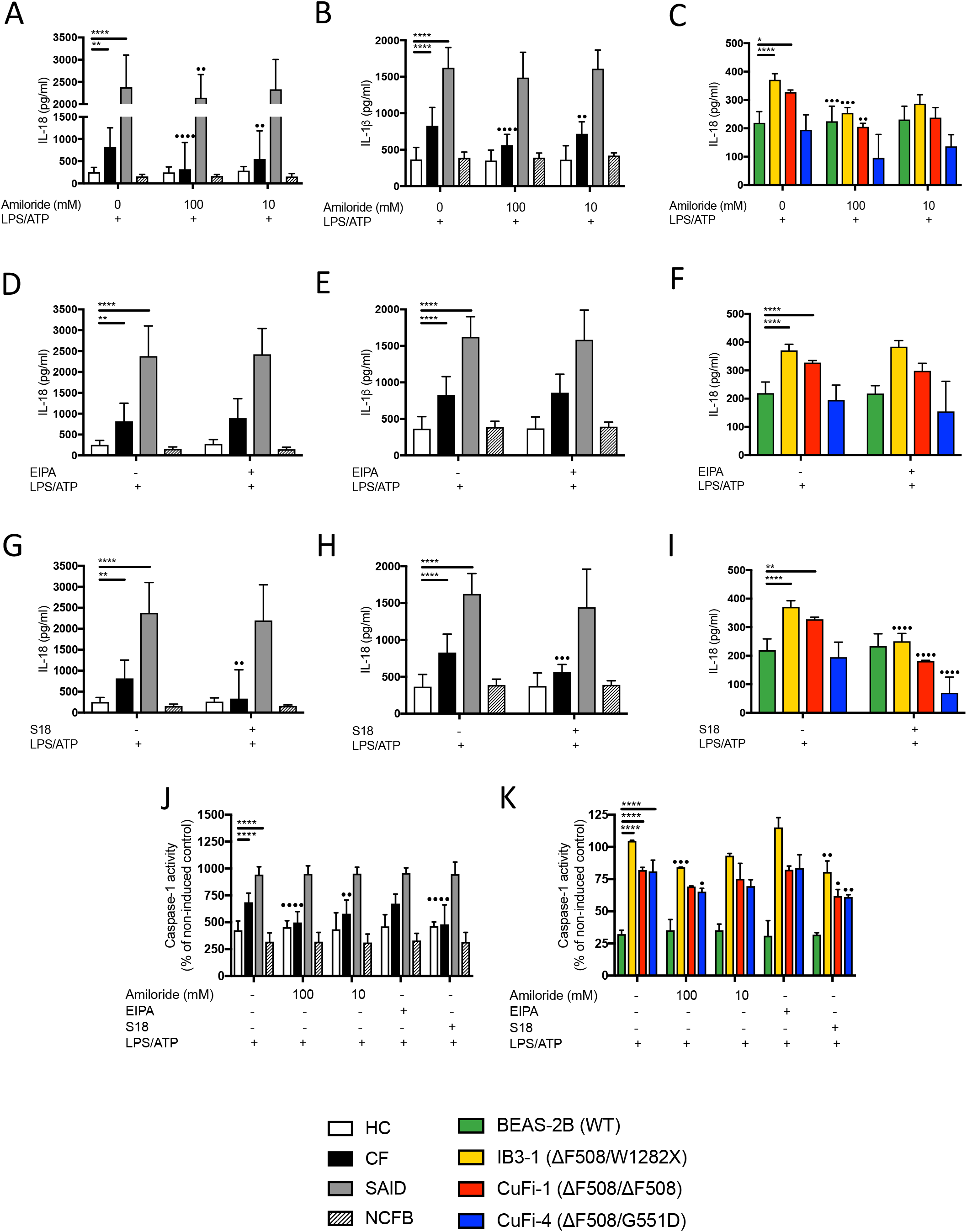
Inhibition of amiloride-sensitive sodium channels modulates inflammation in cells with CF-associated mutations. **A-I** ELISA assays were used to detect IL-18 (A, D, G) and IL-1β (B, E, H) in monocytes from HC (n=10), patients with CF (n=10), SAID (n=4) and NCFB (n=4) and IL-18 in HBECs (n=3) (C, F). Cell stimulation was as follows: Amiloride (100mM or 10mM, 1hour) (A-C), EIPA (10mM, 1 hour) (D-F), S18 derived peptide (25mM, 4 hours) (G-I) were used as a pre-treatment before a stimulation with LPS (10ng/mL, 4 hours) and ATP (5mM) for the final 30 minutes. **J-K** All the above stimulations were used to detect caspase-1 activity using a colorimetric assay in monocytes (J) and HBECs (K). Data Information: A 2-way ANOVA statistical test was performed with Tukey post-hoc correction (p values * = 0.05, **= 0.01, ***= 0.001 and ****= 0.0001).

Our data suggest that dysregulation of amiloride-sensitive Na^+^ channel activity, specifically ENaC in human CF, further potentiates NLRP3 inflammasome activity, via enhanced Na^+^ influx and K^+^ efflux. To our knowledge, this is the first time that the characteristic increase in Na^+^ influx in CF has been associated with K^+^ efflux, upon stimulation.

### Inhibition of amiloride-sensitive sodium channels modulates metabolism in CF

To further examine the extent to which dysregulated Na^+^ flux influences cellular metabolism in patients with CF, and whether this contributes to the observed inflammatory phenotype in CF. A significant amount of cellular energy is utilised to tightly control Na^+^ and K^+^ flux, in order to maintain stable membrane potential and electrochemical equilibrium (Wynne, Zou et al., 2017). A notable result of elevated Na^+^ influx is increased activity of the Na^+^/K^+^ ATPase pump (Peckham, Holland et al., 1997, Stutts, Knowles et al., 1986), which serves to buffer the elevated intracellular Na^+^ concentrations in order to maintain osmotic homeostasis (Munoz-Planillo et al., 2013). A consequent increase in demand for ATP to fuel this pump may underlie the increased metabolic requirements in patients with CF. Previous studies have shown that a switch to glycolysis often occurs in response to acute inflammatory stimuli in myeloid cells, with a consequent inflammatory cytokine secretion profile and macrophage phenotype (Lamkanfi, 2011, Tannahill, Curtis et al., 2013a, Velsor, Kariya et al., 2006, Wen, Ting et al., 2012). In addition, the succinate metabolite, acting as an inflammatory signal to increase pro-IL-1β transcription (Liu, Luc et al., 2016, Stutts et al., 1986), is upregulated during glycolysis.

In order to determine the metabolic state in CF, both monocytes and HBECs were stimulated with LPS, with and without inhibition of amiloride sensitive channels (Fig. 7 A-F). We measured the L-lactate production (Fig. 7 A, D), glucose consumption (Fig. 7 B, E) and succinate accumulation (Fig. 7 C, E) intracellularly, all by-products of glycolysis, we were able to establish whether cells from patients with CF, were significantly more glycolytic than HC (Fig. 7 A-C).

**Figure 7:**
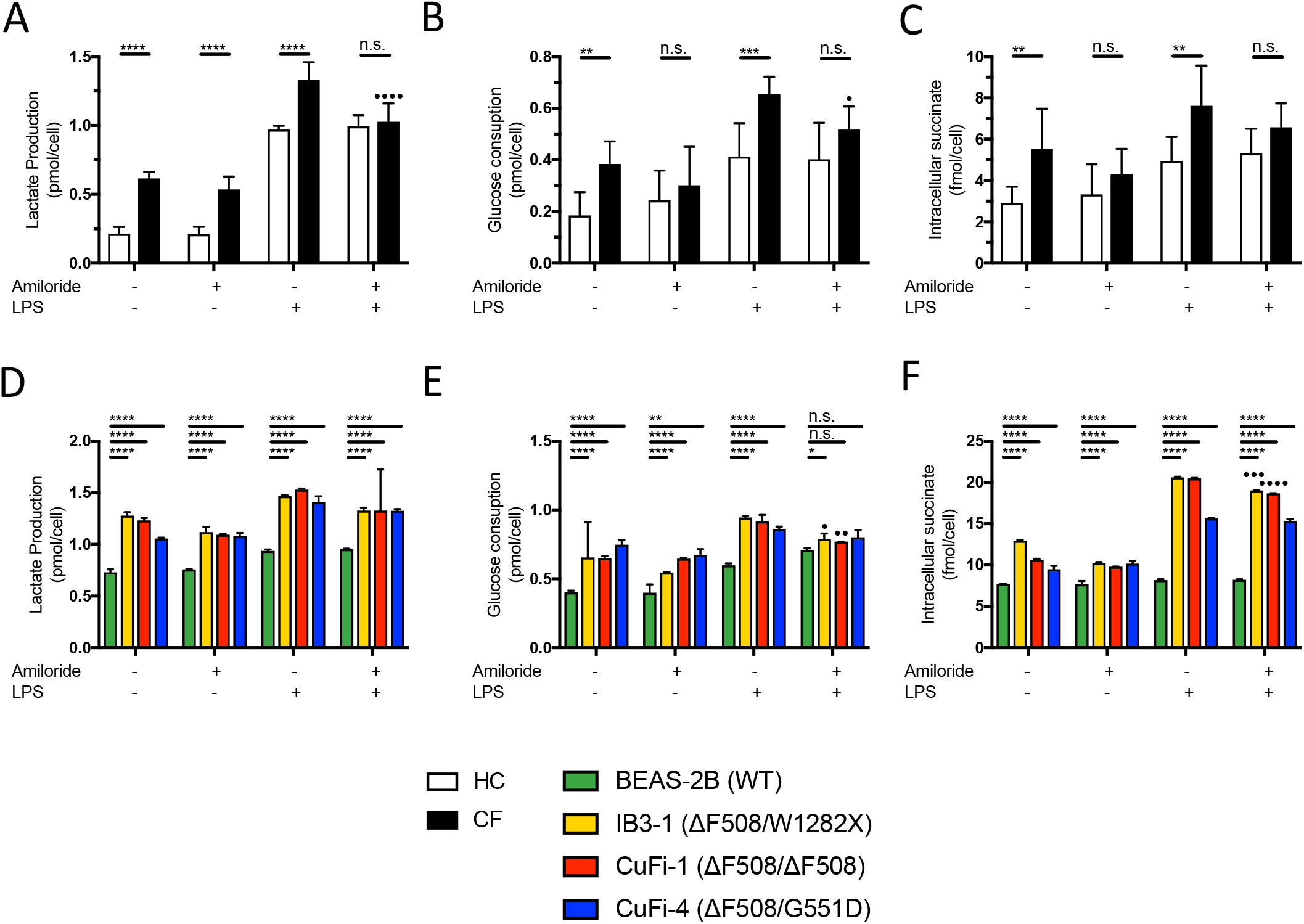
Inhibition of amiloride-sensitive sodium channels modulates metabolism in cells with CF-associated mutations. **A-C** Colorimetric assays were used to detect intracellular L-lactate production (A), glucose consumption (B) and succinate accumulation (C) primary monocytes from HC and CF samples (n=7). Cell were pre-treated with amiloride (100mM, 1 hour) before an LPS (10ng/mL, 4 hours) stimulation. **D-F** Colorimetric assays were used to detect intracellular L-lactate production (D), glucose consumption (E) and succinate accumulation (F) in HBECs (n=3). Cells were pre-treated with amiloride (100mM, 1 hour) before adding LPS (10ng/mL, 4 hours) stimulation. Data Information: A 2-way ANOVA was performed (p values * = 0.05, **= 0.01, ***= 0.001 and ****= 0.0001). (•) indicate significance between treatments within the same cell line (−/+ amiloride).

Patients’ monocytes secreted elevated levels of all of the glycolytic by-products compared to HC, whether unstimulated or following LPS stimulation. Notably, amiloride alone was able to modulate both glucose consumption (Fig. 7 B) and succinate accumulation (Fig. 7 C) in CF monocytes; furthermore in the presence of LPS, amiloride pretreatment significantly reduced lactate production (Fig. 7 A) and glucose consumption (Fig. 7 B), in CF monocytes. All HBEC lines with CF-associated mutations secreted significantly elevated levels of L-lactate (Fig. 7 D), with significant increased glucose consumption (Fig. 7 E) and succinate accumulation (Fig. 7 F), under both unstimulated and LPS-stimulated conditions. Amiloride alone was unable to modulate any of the glycolytic by-products in any of the cell lines tested. However, in the presence of LPS, amiloride decreased glucose consumption (Fig. 7 D) and intracellular succinate (Fig. 7 F) levels in both IB3-1 and CuFi-1 HBEC lines. By inhibiting glycolysis with 2-Deoxy-D-glucose (2-DG), a competitive inhibitor of glucose-6-phosphate, IL-18 and IL-1β secretion in presence of LPS and ATP was inhibited (Supplementary Fig. 2 D-F) whereas TNF levels were not affected (Supplementary Fig. 2 C) (Tannahill et al., 2013a, Tannahill, Curtis et al., 2013b). Together, these data indicate that CF innate immune cells are metabolically reprogrammed, and that the metabolic rate can be reduced by inhibition of overactive Na^+^ flux.

### β-ENaC over-expression in Beas-2B cells increases pro-inflammatory cytokine secretion

In order to recapitulate the ENaC-NLRP3 axis, as revealed by the data in this study, we overexpressed the β-ENaC chain in the WT HBEC line, BEAS-2B. This has previously been explored in a β-ENaC Tg-mouse model of CF, which recreated a CF-like lung disease, with mucous plugging and excessive inflammation (Mall et al., 2004a, Zhou et al., 2011). We studied the intracellular molecular mechanisms that are disrupted by β-ENaC overexpression, to augment our evidence that excessive ENaC-mediated sodium flux modulates the inflammatory phenotype of cells with CF-associated mutations. The β-chain of ENaC (*SCNN1B*) is the rate-limiting chain that completes the formation of a fully functional ENaC channel (Mall et al., 2004a, Zhou et al., 2011). A transient transfection of *SCNN1B* alongside a pcDNA3.1 vector only control for 48 hours increased β-ENaC expression in the BEAS-2B HBEC line (Fig. 8 A); non-transfected cells secreted IL-18 only upon stimulation with both LPS and ATP, whereas cells with overexpression of β-ENaC secreted comparable levels of IL-18 under both unstimulated conditions and LPS alone (Fig. 8 B). Overexpression of β-ENaC induced elevated IL-18 secretion, after LPS and ATP stimulation (Fig. 8 B). These data support the hypothesis that excessive ENaC-mediated sodium influx drives NLRP3 inflammasome activation in CF.

**Figure 8:**
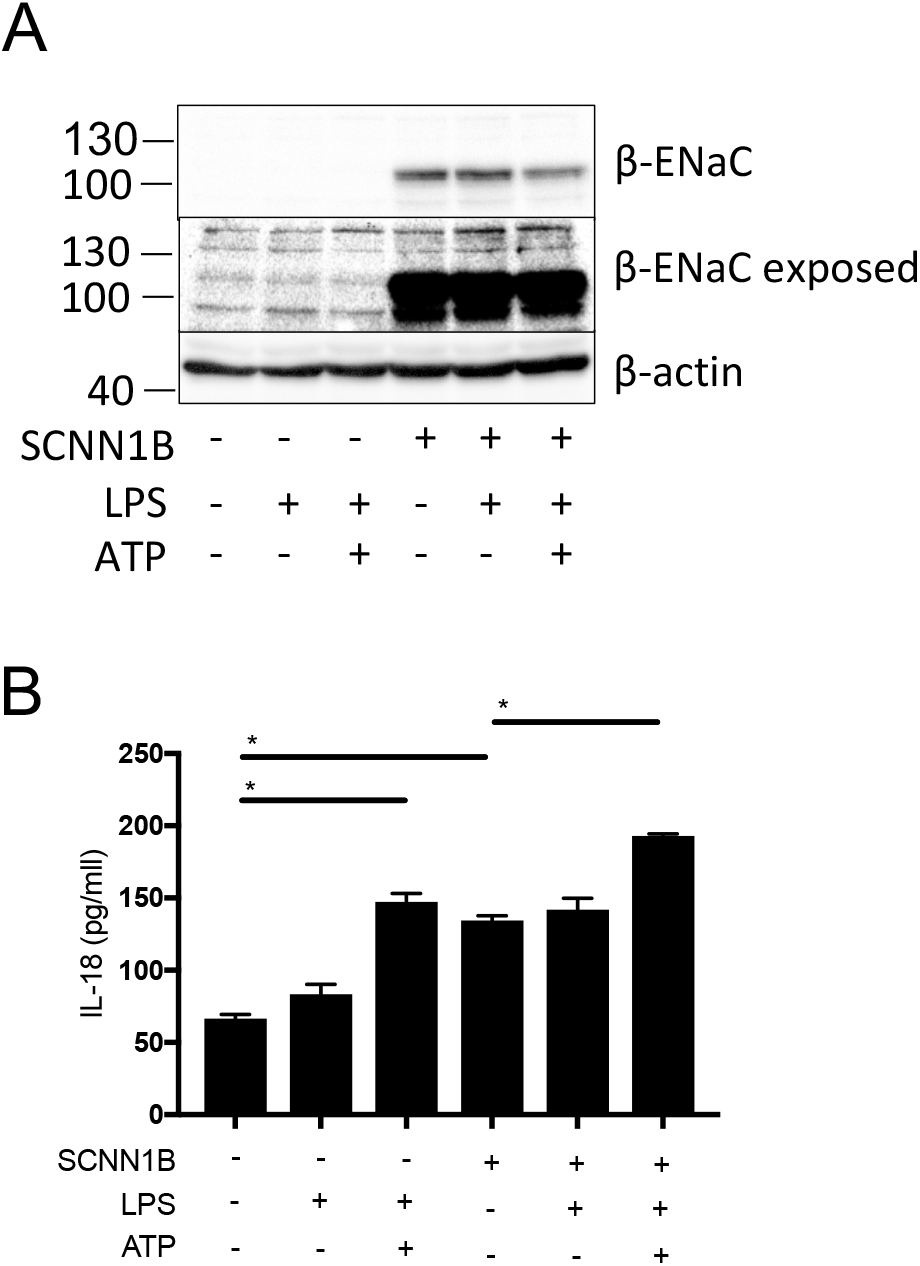
SCNN1B over-expression in BEAS-2B cells increases pro-inflammatory cytokine secretion. **A-B** BEAS-2B cells were transiently transfected with 10*μ*g SCNN1B cDNA (+) or a pcDNA3.1 vector only control (-) for 48h then stimulated with LPS (10ng/mL, 4 hours) and ATP (5mM) for the final 30 minutes (n=3). Cells were lysed and immunoblotted for β-ENaC and β-actin (A). ELISA assays were used to detect IL-18 in the supernatant (B) fraction. Data Information: A 2-way ANOVA statistical test was performed with Tukey post-hoc correction (p values * = 0.05, **= 0.01, ***= 0.001 and ****= 0.0001).

## Discussion

While inflammation in patients with CF is believed to be driven predominantly by bacteria, fungi, and viruses, some CFTR mutant animal models have shown that airway inflammation and bronchiectasis can occur under sterile conditions during infancy (Keiser, Birket et al., 2015) (Rao & Grigg, 2006). The primary consequence of mutations in the CFTR gene is defective CFTR anion transport and increased sodium transport through dysregulation of ENaC. Under *in vivo* conditions, basal transepithelial nasal potential differences in patients with CF are more negative than in HC and show an enhanced depolarising response to amiloride, reflecting an inhibition of excessive ENaC-mediated sodium influx (De Boeck, Derichs et al., 2011). Here we show, for the first time, that increased basal sodium flux is pivotal to the observed increase in potassium efflux and downstream NLRP3 inflammasome activation in both primary monocytes and HBECs of patients with CF.

Autoinflammation is defined as an inappropriate inflammatory response, driven by dysregulated innate immune cells, without any evidence of either antigen-driven T cells, B-cells, or associated autoantibodies (Kastner, Aksentijevich et al., 2010, McDermott & Aksentijevich, 2002, McDermott, Aksentijevich et al., 1999, McGonagle & McDermott, 2006, Pathak, McDermott et al., 2016, Peckham, Scambler et al., 2017, Stoffels & Kastner, 2016). Adaptive immune cells are sometimes recruited in response to the downstream consequences of autoinflammation, with increased susceptibility to infection, and progression autoimmunity and hyperinflammation (Wekell, Karlsson et al., 2016). Based on these data and cited literature presented in this study, CF may be characterised as an autoinflammatory disease, in part driven by aberrant ion flux and recurrent infections. In CFTR knockout ferrets, long term antibiotics have been used to protect the lung from infections. In this model animals still developed inflammation, bronchiectasis and mucus accumulation despite the absence of infection (Rosen et al 2108). Indeed, the NLRP3 inflammasome can be primed by proinflammatory cytokines, such as TNF, although bacterial components provide a far more potent stimulus.

We have examined primary innate immune cells and HBEC lines with CF-associated mutations *in vitro,* and discovered that excessive ENaC-mediated sodium influx in cells with CF-associated mutations prime the NLRP3 inflammasome for excessive and inappropriate IL-18 and IL-1β secretion. We have also shown that the nature of systemic serum cytokine signature from patients with CF is comparable to that of patients diagnosed with SAID, and characterised by release of proinflammatory IL-1-type cytokine family members (IL-1 and IL-18) associated with inflammasome activation. This proinflammatory signature extends to the phenotypes of both monocytes and macrophages, with a marked deficiency of both classical monocytes and M2-type macrophages, and a significantly impaired anti-inflammatory IL-10 response in this condition. Furthermore, both monocytes and HBECs are hyper-metabolic in CF, indicating the induction of an inflammatory effector phenotype. When we explored the type of inflammation involved, we found a hyperresponsive NLRP3 inflammasome, exclusively in cells with CF-associated mutations, which was comparable to monocytes from patients diagnosed with SAID. This NLRP3 inflammasome activation extended beyond IL-18 and IL-1β secretion, with evidence of an increased propensity for pyroptotic cell death and associated release of various NLRP3 inflammasome components, such as ASC, and the induction of IFN-γ secretion in the PBMC population. This type of inflammation has similarities to autoinflammation, particularly the IL-1/IL-18 inflammasome signature observed both in serum and also *in vitro*.

The differential secretion of IL-18 and IL-1β observed in this study is noteworthy, with IL-1β secretion being undetectable in HBEC lines, under the conditions used. This disparity may reflect the function of these two inflammasome-processed zymogens; IL-18 is a cytokine that induces recruitment of neutrophils and Th17 differentiation, as well as IFN-γ secretion, whereas IL-1β is an intrinsically more destructive cytokine, acting systemically to induce fever, proliferation, differentiation, apoptosis and sensitivity to pain. IL-1β secretion is tightly regulated by a highly controlled process, involving its own gene expression as well as inflammasome priming and assembly, whereas IL-18 is constitutively expressed and only depends on the two signals for inflammasome priming and assembly. In addition, IL-1β is rapidly sequestered by IL-1Ra and soluble IL-1RI and –RII, once secreted, which makes IL-1β particularly difficult to detect, and helps explain the severe phenotype of the rare autoinflammatory disease, deficiency of the interleukin-1–receptor antagonist (*DIRA*), caused by homozygous mutations in the IL1RN gene, which encodes IL-1Ra protein (Aksentijevich, Masters et al., 2009).

Targeting inflammation in CF is a much-debated topic, particularly in finding a balance between controlling bronchiectasis on the one hand and losing control of intercurrent infection on the other. The data presented here may suggest that targeting IL-18 may be therapeutically beneficial, by preventing adaptive cell airway infiltration whilst maintaining a potent IL-1β response to infection (Gabay, Fautrel et al., 2018b); however NLRP3 activation has a protective role in animal models of induced colitis (Allen, TeKippe et al., 2010) which may run contrary to the expectation that reduced NLRP3 expression might reduce inflammation in the bowel. Furthermore, IL-18 has an epithelial protective role in promoting the repair of gut epithelium (Zaki, Vogel et al., 2010), and further ex vivo studies and in models of CF are required to fully elucidate the potential benefits or, indeed, hazards of IL-18 blockade (Gabay, Fautrel et al., 2018a). It is notable that inhibition of TLR4 signalling was the most effective means of blocking excessive NLRP3 activation in our study, suggesting that targeting this pathway may be a therapeutic option (avenue) to treat this condition. These novel data, along with the observed elevated K^+^ efflux in cells with CF-associated mutations, indicate that excessive ENaC-mediated sodium influx lowers the threshold for NLRP3 assembly, rather than actually priming this key intracellular component of innate immune defenses.

Based on recent studies on metabolic regulation of immune responses (Ganeshan & Chawla, 2014), including the glycolytic shift in inflammation (Kelly & O’Neill, 2015), we examined the metabolic pathways employed by cells with and without CF-associated mutations. Recent discoveries have delineating cellular metabolism as a marker of inflammation, with a metabolic shift from oxidative phosphorylation to glycolysis, is characteristic of a cell’s requirement to fuel acute effector cell functions, in the presence of an inflammatory drive. Intracellular metabolism in monocyte/macrophages and dendritic cells (DCs) has provided new insights into the functioning of these critical regulators of innate and adaptive immunity (Buck et al., 2017) and highlighted new therapeutic options for the management of CF. Here we have shown that both monocytes and HBEC lines with CF-associated mutations have increased metabolic activity, with induction of glycolysis. Of the glycolytic byproducts elevated in this study, L-lactate and succinate are particularly significant. With CFTR absent or dysfunctional, the loss of HCO3- transport will enable ASL to become acidic. Also, dysregulated metabolism can contribute to the acidic nature of ASL in CF (Garnett et al., 2016). A potential molecular mechanism for a switch to glycolysis in cells with CF-associated mutations may be that the excessive ENaC-mediated sodium influx drives Na^+^/K^+^ATP-gated channel activity to restore ionic homeostasis. As Na^+^/K^+^ATP-gated channel activity is elevated in CF (Peckham et al., 1997) this will increase demand for ATP and encourage a switch to glycolysis, and IL-1β secretion. Future studies will aim to reach an understanding of how ionic fluxes influence immunometabolism and how glycolysis may be targeted with small molecules in patients with CF.

Previous studies have highlighted the clinical importance of ENaC-mediated sodium flux in CF, attributing increased sodium and water absorption and corresponding reduction of ASL volume, to a subsequent reduction in mucus transport and defective pathogen clearance. However, the molecular consequences of excessive ENaC-mediated sodium influx have not been reported in CF; we provide a novel connection between excessive ENaC-mediated sodium influx, a characteristic feature of CF, and NLRP3 inflammasome activation in cells with CF-associated mutations. Through inhibition of amiloride-sensitive sodium channels with small molecules and peptides, we were able to reduce the excessive NLRP3 inflammasome activation, reverse the switch to glycolysis and associated IL-18 and IL-1β secretion *in vitro*. These novel findings highlight the importance of excessive ENaC-mediated sodium influx as a disease mechanism, as well as highlighting its potential as a therapeutic target in patients with CF. Targeting amiloride-sensitive sodium channels, such as ENaC, has been previously attempted, using amiloride in the early 1990s, with little efficacy, due to amiloride’s short half-life and limited effectiveness (Graham, Hasani et al., 1993, Knowles, Church et al., 1990). There are more recent studies showing that pre-emptive administration of amiloride may be more effective, based on studies with the βENaC-Tg mouse (Zhou et al., 2008). Developing new ENaC inhibitors is an ongoing mission of multiple academic and pharmaceutical groups: high-throughput screening of small molecules as well as small interfering RNAs (siRNAs) (Gianotti, Melani et al., 2013), targeting the proteases that activate ENaC (Gaillard, Kota et al., 2010) and modifying endogenous ENaC inhibitors to increase their viability in the acidic CF lung (Garcia-Caballero, Rasmussen et al., 2009), are all potentially useful approaches for *in vivo* ENaC inhibition in patients with CF. Therefore, inhibitors of amiloride-sensitive sodium channels or stimulation of alternative Cl^−^ channels (Domingo-Fernández, Coll et al., 2017) are two therapeutic strategies that may modulate downstream NLRP3 inflammasome activation regardless of CFTR genotype, and potentially reduce the inflammatory burden in CF.

We have found that excessive ENaC-mediated sodium influx lowers the threshold for potassium efflux-dependent NLRP3 inflammasome activation in cells with CF-associated mutations. Hypersensitive NLRP3 inflammasome activation in CF induces proinflammatory serum and cellular profiles, particularly involving IL-18 release from HBECs, that may be abrogated by inhibition of amiloride-sensitive sodium channels. Our study challenges current ideas on the primary molecular defects in CF and proposes that therapies targeted against the ENaC-NLRP3 axis will be beneficial in controlling inappropriate autoinflammatory responses in CF. It furthermore supports the concept that pulmonary inflammation, under some circumstances, is disassociated from infection in CF, and these patients may exhibit a chronic underlying autoinflammatory state with infection-related exacerbations. These findings have major implications for potential therapeutic intervention in CF, and all dysrergulated pathways that underlie this autoinflammatory state, need to be corrected for successful resolution of the pathology of CF.

## Materials and Methods

### Patients’ samples

Patients with cystic fibrosis (CF) were recruited from the Adult Cystic Fibrosis Unit at St James’s University Hospital, Leeds. All patients had two disease-causing CFTR mutations and clinical features consistent with the diagnosis of CF (F508del/F508del, n = 35; F508del/c.1521_1523delCTT, n =2; 3849+10KB C>T/F508del, n=1; c.1040G>C/R347P, n=1; F508del/621+1 (G>T), n=1). Patients included in Figs 2-8 were all F508del/F508del with no sign of infection. Patients with NCFB and SAID were recruited from the Leeds Regional bronchiectasis unit and Department of Clinical Immunology and Allergy, respectively. All patients with a SAID had characterized mutations in a known disease-causing gene (TRAPS n=2, Muckle-Wells n=2, A20 haploinsufficiency n=1, PAAND n=1, MEFV n=2, FMF n=2, HIDS n=2 and Schnitzler syndrome n=1) (Table 1). Age and sex matched healthy controls were recruited from the research laboratories at the Wellcome Trust Benner Building, St James’s University Hospital, Leeds, UK. All patient samples collected are approved by the Health Research Authority REC reference 17/YH/0084.

**Table 1.**
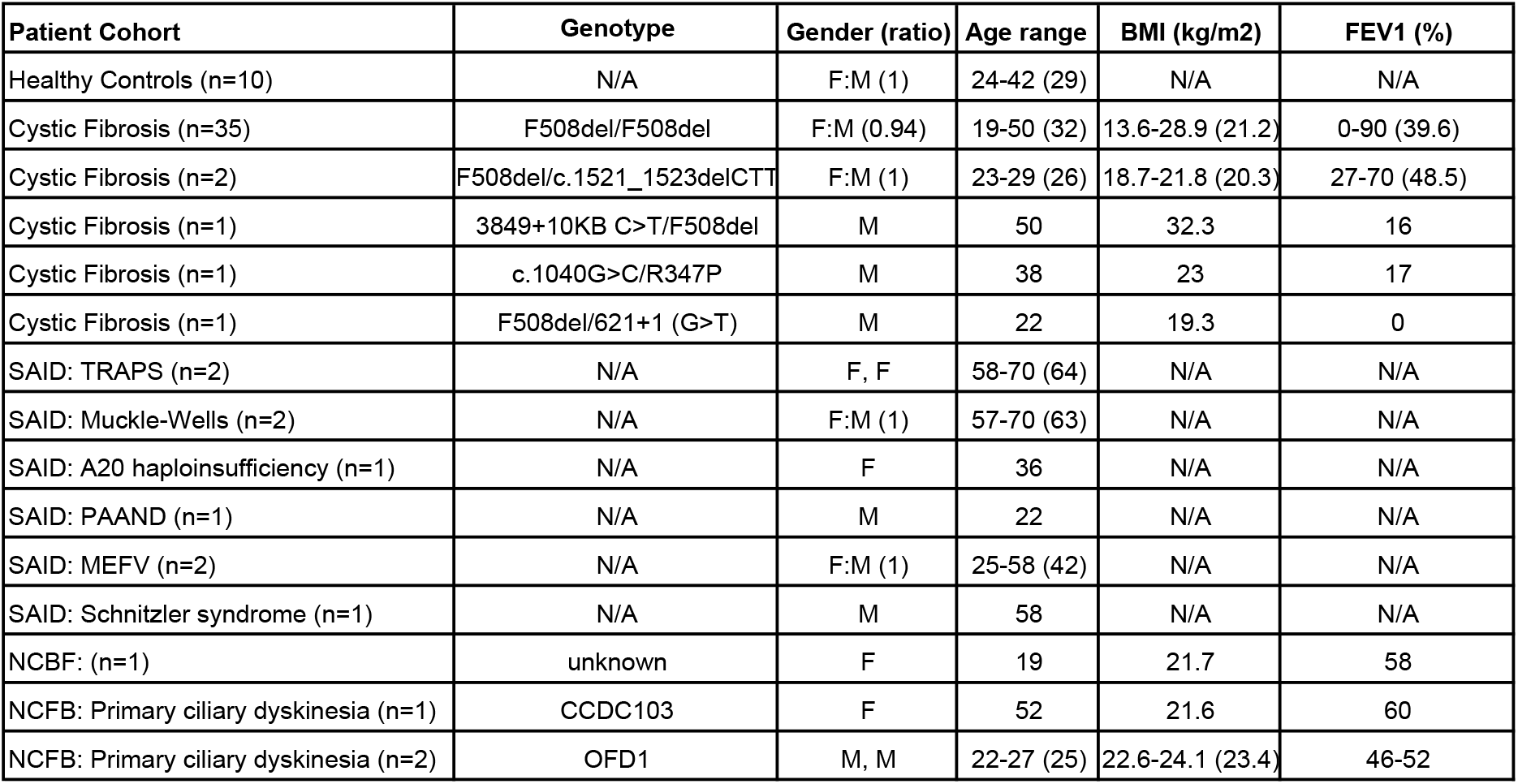
Clinical Data

| Patient Cohort | Genotype | Gender (ratio) | Age range | BMI (kg/m2) | FEV1 (%) |
| --- | --- | --- | --- | --- | --- |
| Healthy Controls (n=10) | N/A | F:M (1) | 24-42 (29) | N/A | N/A |
| Cystic Fibrosis (n=35) | F508del/F508del | F:M (0.94) | 19-50 (32) | 13.6-28.9 (21.2) | 0-90 (39.6) |
| Cystic Fibrosis (n=2) | F508del/c.1521_1523delCTT | F:M (1) | 23-29 (26) | 18.7-21.8 (20.3) | 27-70 (48.5) |
| Cystic Fibrosis (n=1) | 3849+10KB C>T/F508del | M | 50 | 32.3 | 16 |
| Cystic Fibrosis (n=1) | c.1040G>C/R347P | M | 38 | 23 | 17 |
| Cystic Fibrosis (n=1) | F508del/621+1 (G>T) | M | 22 | 19.3 | 0 |
| SAID: TRAPS (n=2) | N/A | F, F | 58-70 (64) | N/A | N/A |
| SAID: Muckle-Wells (n=2) | N/A | F:M (1) | 57-70 (63) | N/A | N/A |
| SAID: A20 haploinsufficiency (n=1) | N/A | F | 36 | N/A | N/A |
| SAID: PAAND (n=1) | N/A | M | 22 | N/A | N/A |
| SAID: MEFV (n=2) | N/A | F:M (1) | 25-58 (42) | N/A | N/A |
| SAID: Schnitzler syndrome (n=1) | N/A | M | 58 | N/A | N/A |
| NCFB: (n=1) | unknown | F | 19 | 21.7 | 58 |
| NCFB: Primary ciliary dyskinesia (n=1) | CCDC103 | F | 52 | 21.6 | 60 |
| NCFB: Primary ciliary dyskinesia (n=2) | OFD1 | M, M | 22-27 (25) | 22.6-24.1 (23.4) | 46-52 |

### Sample preparation

Peripheral blood mononuclear cells (PBCs) were isolated from whole blood using the density gradient centrifugation. Whole blood were mixed with equal volume of PBS, carefully layered onto of Lymphoprep® (Axis-Shield) and centrifuged at 1100*xg* for 20 min without brakes. The white buffy layer is removed and washed twice in PBS by centrifuging at 1100*xg* for 10 min. PBMC pellet is resuspended in complete RPMI medium (RMPI medium containing 10% heat inactivated foetal bovine serum, 50 U/ml penicillin, 50 μg/ml streptomycin).

Monocytes were isolated by negative selection from PBMCs using the monocyte isolation kit II (Miltenyi Biotec). Pelleted PBMCs were resuspended in 30 μl of buffer (autoMACS Rinsing Solution containing 0.5% BSA). This was mixed with 10 μl FcR Blocking Reagent followed by 10μl of biotin-conjugated antibodies and incubated at 4°C for 10 min. Next 30 μl of buffer were added together with 20 μl of anti-biotin microbeads and incubated for an additional 15 min at 4°C. This whole mixture was washed with 2 ml of buffer and centrifuged at 300 *xg* for 10 min. The cell pellet was resuspended in 500 μl of buffer and past down a MS column (Miltenyi Biotec) on a magnetic stand.

### Cell culture

Human cell lines BEAS-2B, CuFi-1, CuFi-4 and IB3-1 were purchase from ATCC (UK). BEAS-2B and IB3-1 were cultured in LHC basal medium (Thermo Fisher Scientific) supplemented with 10% FBS, 50 U/ml penicillin and 50 μg/ml streptomycin). CuFi-1 and CuFi-4 were grown on Cell+ surface plates or flasks (Sarstedt) with LHC-9 medium (Thermo Fisher Scientific).

### Cell stimulations

PBMCs at 2×10^6^ cell/ml/well were incubated in 1 ml of complete RPMI medium of a 6-well plate. Cell lines were seeded at 1×10^6^ cells/ml/well cultured in 1 ml of appropriate growth media (see cell culture). All cells were pre-treated with the following compounds where indicated prior to NLRP3 stimulation; Amiloride (100mM, Cayman Chemical) for 1 h, MCC950 (15nM, Cayman Chemical) for 1 h, 2-DG (0.5mM, Agilent) for 2 h, EIPA (10μM, Cayman Chemical) for 1 h, NPPB (100μM, Cayman Chemical) for 1 h, ouabain (100nM, Cayman Chemical) for 24 h, digoxin (100nM, Cayman Chemical) for 24 h, OxPAC (30μg/mL, Invivogen) for 1 h, Ac-YVADD (2μg/mL, Invivogen) for 1 h and CFTR172 (10μg/mL, Cayman Chemical) for 1 h.

Inflammasome stimulation was achieved using either LPS (10ng/mL, Ultrapure EK, Invivogen) for 4 h with the addition of ATP (5mM, Invivogen) at the final 30 min of stimulation, poly(dA:dT) dsDNA (1μg/mL with Lipofectamine 2000, Invivogen) for 1 hour, TcdB (10ng/mL, Cayman Chemical) for 1 h or flagellin (10ng/mL with Lipofectamine 2000, Invivogen) for 1 h in the hour of the LPS stimulation. All incubations were done in a humidified incubator at 37°C, 5% CO_2_.

### ELISA

Cytokines from patient sera and cell cultured media were detected by ELISAs (Human IL-1β CytoSet^TM^, human IL-18 matched antibody pairs, human IL-1Ra CytoSet^TM^, human TNF-α antibody pair and human IL-6 CytoSet^TM^) (Invitrogen), as per the manufactures recommendations. In general, ELISA plates were coated with 100 μl cytokine capture antibody in PBS overnight at 4°C. The plates were washed 3 times with PBST (PBS containing 0.5% Tween 20) and the wells blocked in 300 μl assay buffer (0.5% BSA, 0.1% Tween 20 in PBS) by incubating for 1 h. The plates were washed twice with PBST and 100 μl of sera/culture supernatants, together with appropriate standards, were added to wells in duplicates. Immediately 50 μl of detection antibody were added to all wells and incubated for 2 h. After the incubation the plates were washed 5 times with PBST and 100 μl of tetramethybenzidine (TMB) substrate solution (Sigma) were added to all wells and incubated for 30 min. Colour development was stopped by adding 100 μl of 1.8N H_2_SO_4_. And absorbance measured at 450 nm and reference at 620 nm. Note all incubation steps were done at room temperature with continual shaking at 700 rpm.

### RT-qPCR

Cells were washed in PBS, pelleted and immediately lysed in 1 ml TRIzol™ Reagent (Ambion Life technologies) and RNA extracted using the PureLink RNA mini kit (Ambion). Chloroform (200 μl) were added to each sample and mixed vigorously for 15 sec and left to stand for 2 min at room temperature. These were centrifuged at 12000 *xg* for 15 min at 4°C. The top clear phase was transferred to a fresh tube and mixed with equal volume of 70% ethanol. This mixture was transferred to a spin cartridge, with a collection tube, and centrifuged at 12000 *xg* for 15 sec at room temperature. The waste was disposed of and the spin cartridge was centrifuged one more time. The spin cartridge was washed 700 μl of wash buffer I and centrifuged at 12000 *xg* for 15 sec. A second wash with 500 μl wash buffer II (containing ethanol) were added to the spin cartridge and centrifuged at 12000 *xg* for 15 sec followed by further spin for 1 min. RNA was recovered by adding 30 μl of RNase free water to the spin cartridge, incubated for 1 min and centrifuged for 2 min.

The High Capacity cDNA Reverse Transcription kit (Applied Biosystems) was used to convert the RNA to cDNA according to the manufacturer’s instructions. SYBR Green PCR Master mix (Applied Biosystems) was used to analyse the expression of selected primers in the cDNA. SYBR Green master mix (24 μl) and specific forward and reverse primers (300 nM) to the genes of interest. This mixture was loaded onto a 96 well PCR plate and loaded on to the PCR machine 7500 Real-Time PCR System (Applied Biosystems) programmed with the following cycling conditions: 1 cycle at 50°C for 2 min, 1 cycle at 95°C for 10 min, 50 cycles at 95°C 15 sec, 60°C 1 min, 95°C 15 sec, 1 cycle at 60°C for 20 sec, 1 cycle at 95°C for 15 sec and 1 cycle at 60°C for 1 min (Mathews *et al*. 2014). TaqMan assays were done in the QuantStudio 5 Real-Time PCR instrument (ThermoFisher Scientific).

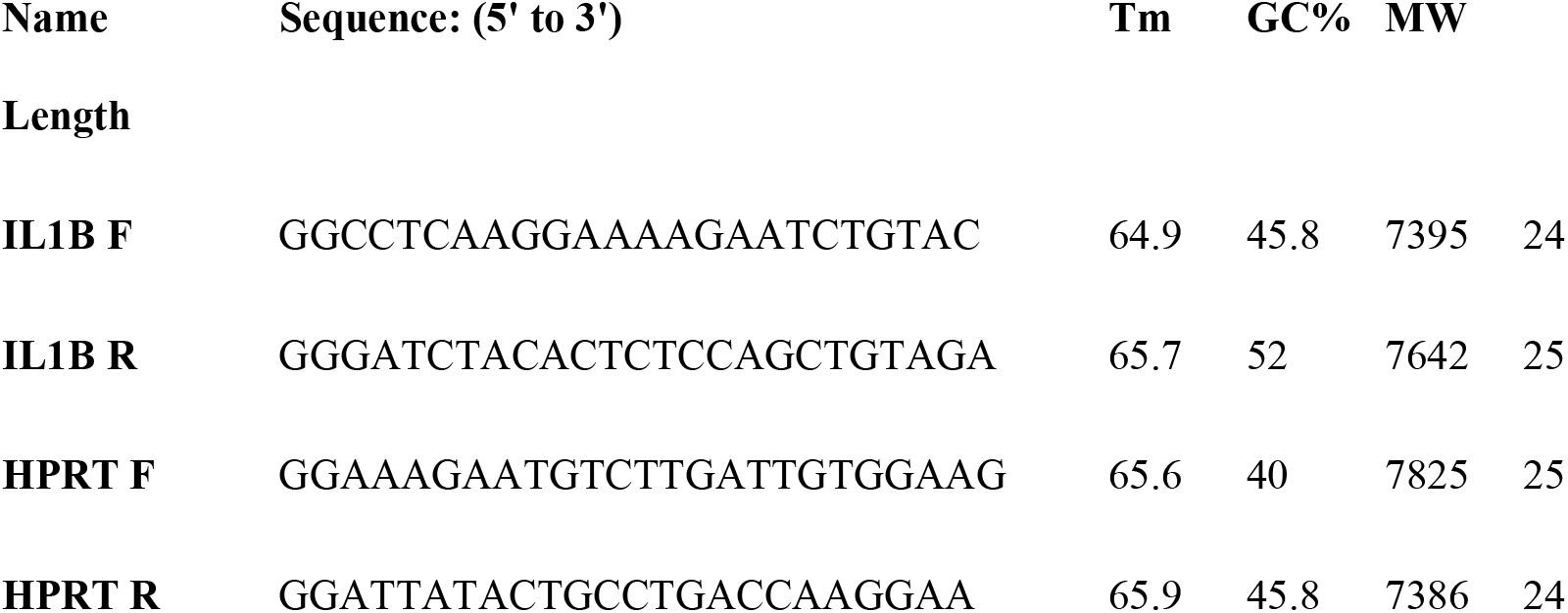

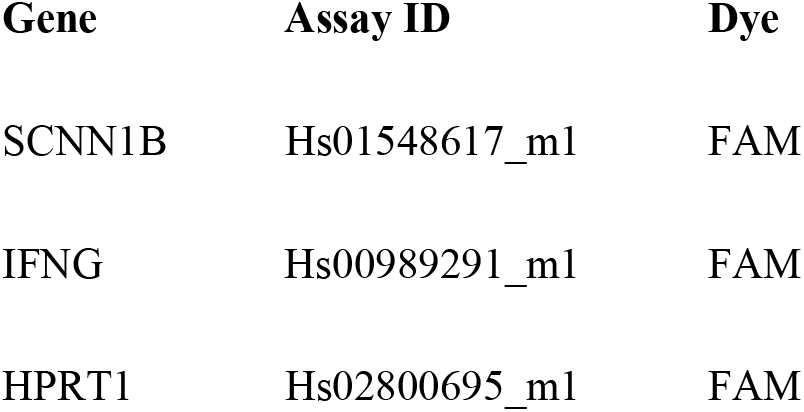
The Taqman® primers used in this study are detailed below:

The Taqman® primers used in this study are detailed below:
| Gene | Assay ID | Dye |
| --- | --- | --- |
| SCNN1B | Hs01548617_m1 | FAM |
| IFNG | Hs00989291_m1 | FAM |
| HPRT1 | Hs02800695_m1 | FAM |

### Western Blotting

Samples were made up in dissociation buffer [1x dissociation buffer (100mM Tris-HCl, 2% (w/v) sodium dodecyl sulfate, 10% (v/v) glycerol, 100mM dithiothreitol, 0.02% (w/v) bromophenol blue, pH 6.8] and heated at 95°C for 5 min. Protein concentrations were determined using the bicinchoninic acid (BCA) assay (Pierce). Equal proteins concentration were load and resolved by 10% SDS-PAGE on Tris-glycine gels and then transferred to Hybond PVDF membranes (GE Healthcare). Following electrotransfer in Towbin buffer (25 mM Tris, 192 mM glycine, pH 8.3, 20% methanol) at 100 V for 1 h, the membranes were blocked for 1 h in blocking solution (PBS containing 0.1% Tween 20 and 5% (w/v) non-fat milk). After 3 washes in PBST (PBS with 0.5% Tween 20), primary antibodies were incubated with PVDF membrane overnight at 4°C. The membrane was washed 3x with PBST and secondary antibody-HRP conjugate were added and incubated for 2 h with constant rocking at room temperature. The membrane was washed 5x with PBST and 3 ml of ECL® detection system (Immobilon chemiluminescent HRP substrate, Millipore) were onto the membrane for 5 minutes, before being imaged with the ChemiDoc Imaging system (Bio-Rad). Primary antibodies used: rabbit anti-SCNN1B (Avia Systems Biology; 1/500 dilution), rabbit anti-actin β (GeneTex) at 1/20000 dilution. Secondary antibodies used: anti-rabbit/mouse IgG horseradish peroxidase-conjugated (Cell Signalling) were diluted at 1/4000. All antibodies were diluted in PBS containing 0.1% Tween 20 and 2% BSA.

### Capsase-1 activity

A colorimetric assay (Caspase-1 Colorimetrix Assay, R&D Systems) was to determine caspase-1 activity via cleavage of a caspase-specific peptide conjugated to a color reporter molecule p-nitroalinine (pNA). The assay was performed in protein lysates and serum. Protein concentrations in the lysate were determined by BCA assay (Pierce).

### Transient Transfection

BEAS-2B cells were transiently transfected with 10μg of SCNN1B cDNA (Addgene) or pcDNA3.1 vector only control (gift from Professor N.M Hooper, Manchester University) using Lipofectamine 2000 reagent (Thermo Fisher Scientific) for 48h as per manufacturer’s instructions. Cells were harvested and Western blotted for SCNN1B to confirm expression.

### Lactate dehydrogenase (LDH) Cytotoxicity Assay

The LDH Cytotoxicity Assay Kit (Pierce) was used, as per the manufacturer’s recommendations, with and without caspase-1 inhibitor, Ac-YVAD-cmk (InvivoGen), as a control for inflammasome dependent cell death (pyroptosis). Optimal cell density was calculated, and the chemical-mediated cytotoxicity assay was performed, whereby spontaneous and maximal LDH activity were calculated as controls for stimulation-mediated cytotoxicity. Absorbance was measured at 490 nm and 680 nm and subtracted from each other.

### M1 and M2 differentiation and activation

Monocytes were isolated from PBMCs by negative selection using the Monocytes isolation kit (Miltenyi Biotech). Monocytes were seeded at 2×10^6^ cells/mL/well of a 6-well plate and cultured in complete RPMI 1640 supplemented with either 20ng/mL human GM- CSF (PeproTech EC Ltd) for M1 differentiation or 20ng/mL human M-CSF (PeproTech EC Ltd) for M2 differentiation and incubated for 9 days. For M1 activation cells are then stimulated with 100ng/mL human IFN-γ (PeproTech EC Ltd), 100ng/mL TNF and 50ng/mL LPS or for M2 activation, 20ng/mL IL-13 and 20ng/mL IL-4 and incubated for 24 hours (Italiani & Boraschi, 2014, Mia et al., 2014).

### M1 and M2 Macrophage characterization

Differentiated and activated M1 and M2 macrophages were characterized by surface marker expression and detection of intracellular cytokines by flow cytometry. Surface markers for M1-type (CD14+ CD16+ HLA-DR+ CD274+ CD86+) and M2-type (CD14+ CD16+ CD206+). Intracellular markers for M1-type (TNFHI) and M2-type (IL-10HI). Monocytes were characterized as classical (CD14++CD16−), non-classical (CD14dimCD16++), and intermediate (CD14++CD16+). All antibodies were from BD Biosciences.

### Fluorometric determination of Na+ and K+ concentration

Sodium and potassium sensitive dyes, SBFI and PBFI, respectively (Molecular Probes) were used, as per the manufacturer’s instructions.

### ASC protein aggregates (specks)

Methodology previously published in (Rowczenio, Pathak et al., 2018)

### Cellular metabolism

Live analysis of ECAR and OCR using the cell energy phenotype kit was performed, using the Seahorse XF-96 Extracellular Flux Analyzer (Agilent Seahorse Bioscience), as per the manufacturer’s instructions. ATP, succinate, extracellular glucose and L-lactate (Assay kits, Abcam) were measured, as per the manufacturer’s instructions.

### Statistics

No statistical methods were used to predetermine sample size. All analyses were performed using GraphPad Prism v 7. Differences were considered significant when P < 0.05. All bar graphs were expressed as medium 95% confidence intervals. The Mann-Whitney non-parametric test was performed (p values * = ≤0.05, **= ≤0.01, ***= ≤0.001 and ****= ≤0.0001) when comparing non-parametric populations. A 2-way ANOVA Kruskal–Wallis statistical test with Bennett’s post-hoc analysis was performed when calculating variance between samples (p values * = ≤0.05, **= ≤0.01, ***= ≤0.001 and ****= ≤0.0001).

## Acknowledgements

The authors would like to thank Professor Eric Blair (Leeds), who provided the IB3-1 cell line as a generous gift and Clive Carter (Leeds) for his invaluable laboratory expertise. The authors would also like to thank all the patients and research nurses, particularly Lindsey Gillgrass and Anne Wood, of the Adult Cystic Fibrosis Unit at St. James Hospital, Leeds. A special acknowledgement for Professor Philip Hopkins (Leeds) for instrumental strategic support. The authors would also like to thank Professor Graham Cook (Leeds) and Devon Scambler (Sheffield) for their editorial advice.

## Author contributions

Conception and design: TS, SS, DP, MFM; Acquisition, analysis and interpretation of data: TS, HJG, CW, SP, SLR, JH; drafting the manuscript for important intellectual content: TS, CW, HJG, FM, SS, DP, MFM

## The paper explained

### Problem

Cystic fibrosis (CF) is one of the most common life limiting autosomal recessive diseases affecting Caucasians. It is a multisystem disease affecting many organs and is characterised by recurrent respiratory tract infections, lung damage and progressive respiratory failure. Despite significant improvements in therapy, quality of life and average life expectancy remain reduced for people with CF. Novel approaches to treatment based on improved understanding of the pathogenesis of CF are necessary to improve patient outcomes in this condition. There is evidence that diseases chronicity involves a vicious cycle of mucostasis and infection on a background of constitutive autoinflammation. We used a multipronged immunometabolic approach to decipher new components of the dysregulated network underlying CF.

### Results

We show that NLRP3 inflammasome activation in immune cells underlies the proinflammatory IL-1 family cytokine (IL-1β and IL-18) signature present in patients with CF. Both primary monocytes and human bronchial epithelial cell (HBEC) lines, with CF-associated mutations, hyper-responded to NLRP3 inflammasome stimulation, compared with controls. Increased IL-18 secretion, by both monocytes and HBEC, was particularly significant, with levels comparable to those of patients with systemic autoinflammatory disease (SAID). This hyperresponsiveness was associated with increased intracellular sodium influx, increased potassium efflux, M2-type macrophage deficiency and a shift to glycolytic metabolism which was abrogated by the addition of NLRP3 inhibitors and inhibition of overactive amiloride-sensitive sodium channels, by either amiloride or S18 peptide. Finally, overexpression of the epithelial sodium channel, β-ENaC, increased IL-18 production in control HBECs without CFTR mutations, thereby supporting the notion that excessive ENaC-mediated sodium influx drives NLRP3 inflammasome activation in these patients.

### Impact

The study shows, for the first time, that excess sodium transport in patients with CF regulates inflammation by activating the NLRP3 inflammasome. Sodium channels are a potential therapeutic target to reduce inflammation in CF, either as monotherapy or in combination with small molecule agents that target defects in the CFTR. Overexpression of β-ENaC, in the absence of CFTR dysfunction, increased proinflammatory cytokine secretion and HBECS with CF-associated mutations specifically produce IL-18, suggesting that modulation of this cytokine may present another therapeutic option in this debilitating condition.

---

**Supplementary Figure 1:**
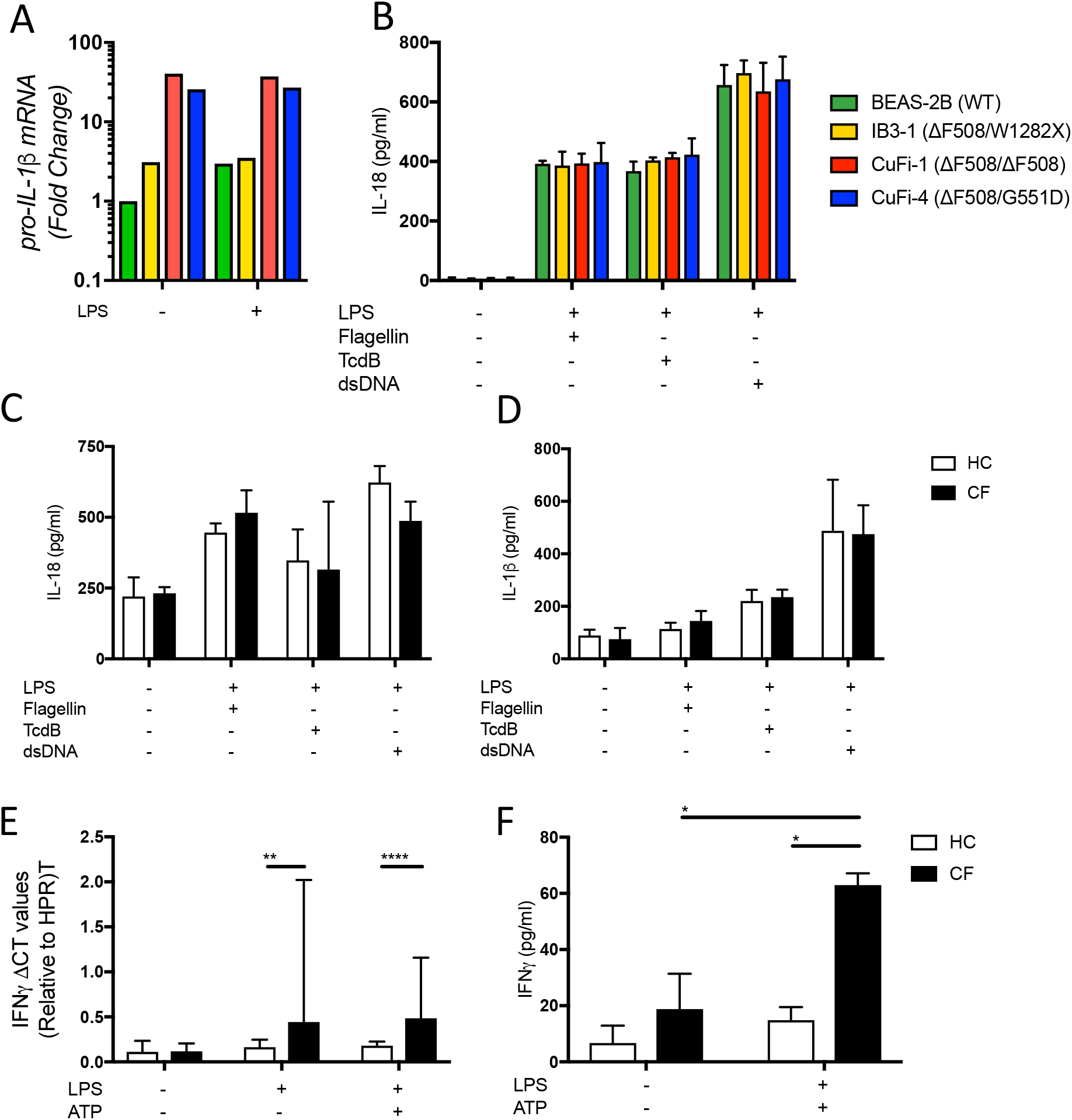
*in vitro* NLRC4, AIM2 and Pyrin Inflammasome activation. **A** Gene expression of pro-IL-1b HBECs (n=1), represented as fold-change. **B-D** ELISA assays were used to detect IL-18 from supernatants of HBECs (A) (n=4) and (C) monocytes (n=10), as well as IL-1b from monocytes (D) (n=10). Cells were stimulated with LPS (10ng/mL) for 4 hours before being stimulated for 4 hours with Flagellin (10ng/mL with Lipofectamine 2000) for NLRC4 inflammasome, TcdB (10ng/mL) for Pyrin inflammasome or poly(dA:dT) dsDNA (1mg/mL with Lipofectamine 2000) for AIM2 inflammasome. **E-F** Taqman® RT-qPCR was used to measure *IFN*g gene expression (A) and Luminex was used to measure IFNg secretion (B) from peripheral blood mononuclear cells (PBMC) populations from HCs and patients with CF-associated mutations (n=10). Data Information: A 2-way ANOVA statistical test was performed (p values * = 0.05, **= 0.01, ***= 0.001 and ****= 0.0001).

**Supplementary Figure 2:**
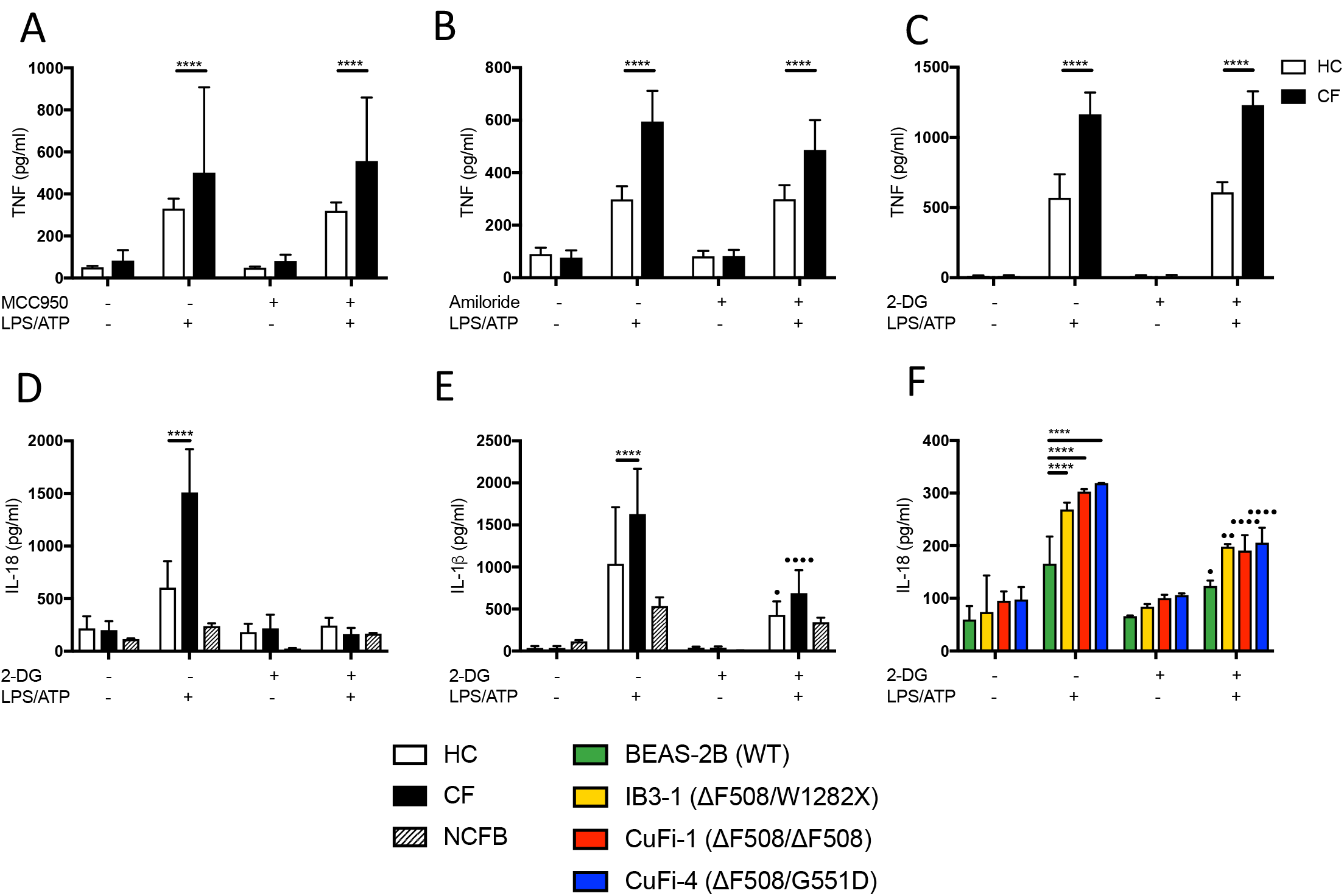
TNF, IL-18 and IL-1b secretion after inhibition with 2-DG, amiloride and MCC950. **A-C** Primary monocytes from HC (n=6) and CF (n=6) were pre-incubated with MCC950 (15mM) (A), 2-DG glycolysis inhibitor (B) and amiloride (C) followed by a stimulation with LPS (10ng/mL, 4 hours), and ATP (5mM) for the final 30 minutes. ELISA assays were used to detect TNF (A-C) in supernatants. **D-F** Primary monocytes from HC (n=6), CF (n=6) and NCFB (n=4) and HBECs (n=4) were pre-incubated with 2-DG glycolysis inhibitor followed by a stimulation with LPS (10ng/mL, 4 hours), and ATP (5mM) for the final 30 minutes. ELISA assays were used to detect IL-18 and IL-1b from monocyte supernatants (D, E) and IL-18 from supernatants of HBECs (F) (n=4). Data Information: A 2-way ANOVA statistical test was performed (p values * = 0.05, **= 0.01, ***= 0.001 and ****= 0.0001).

**Supplementary Figure 3:**
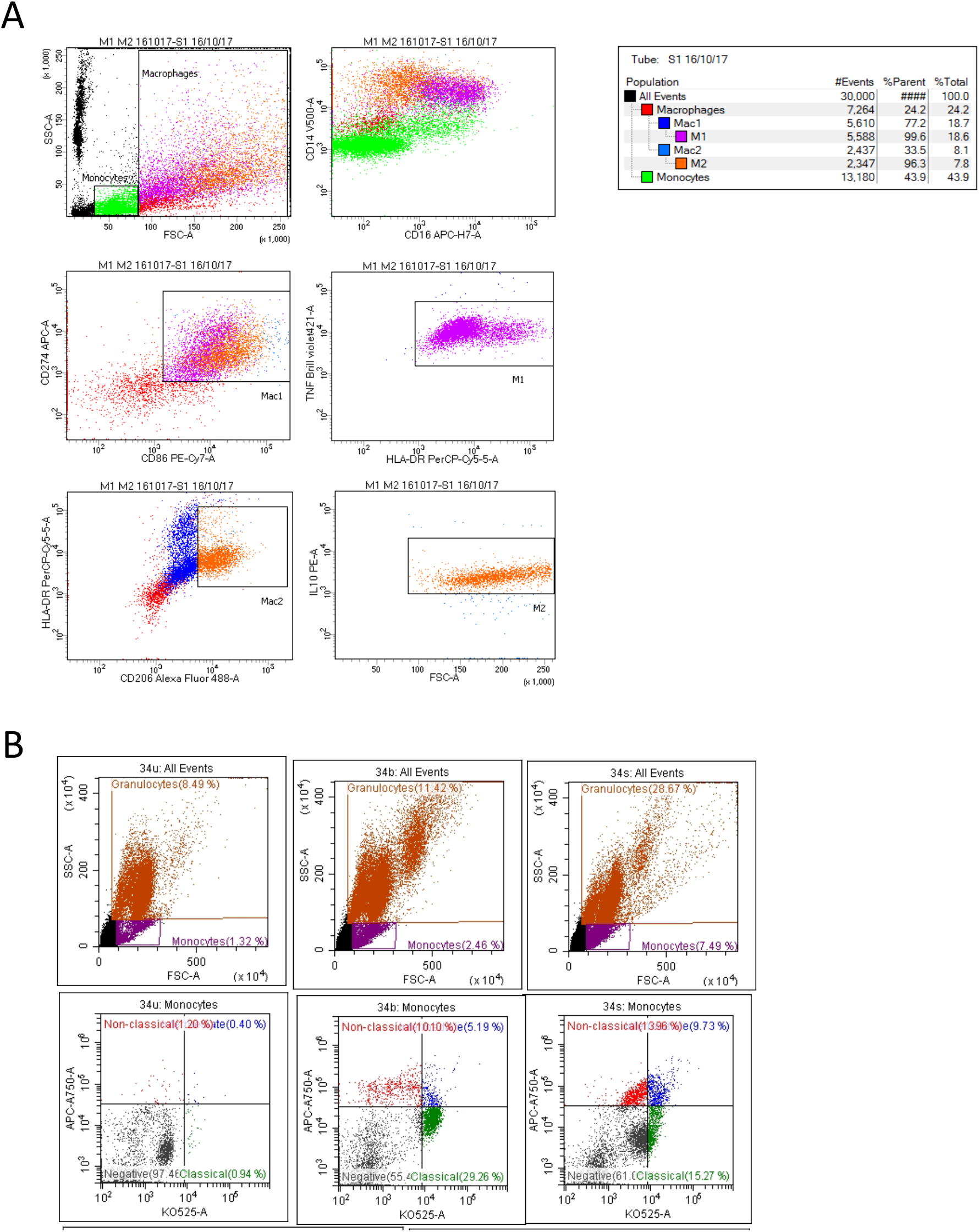
Gating strategy for monocyte and macrophage characterisation. **A** Monocytes from whole blood were differentiated into macrophages and gated based on forward and side scatter. M1-type (markers-CD14^+^ CD16^+^ HLA-DR^+^ CD274^+^ CD86^+^ TNF^HI^). M2-type (markers-CD14^+^ CD16^+^ CD206^+^ IL-10^HI^) (A). **B** Monocytes were gated on based on forward and side scatter and then based on CD14 and CD16 expression as follows: classical (CD14++CD16−), non-classical (CD14dimCD16++), and intermediate (CD14++CD16+) (B).

**Supplementary Figure 4:**
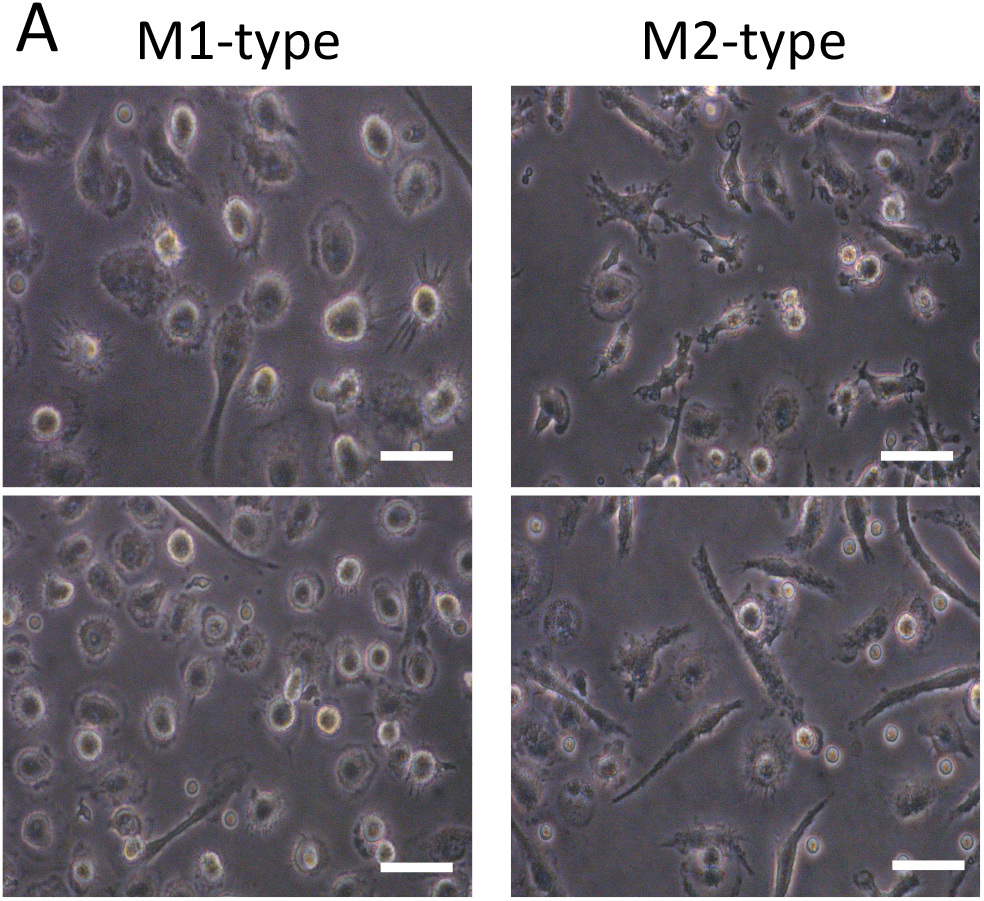
Macrophage morphology. Surface markers for M1-type (CD14+ CD16+ HLA-DR+ CD274+ CD86+) and M2-type (CD14+ CD16+ CD206+) were analyzed using flow cytometry. M1-type macrophage activation was achieved by supplementing growth media with 100ng/mL human IFN-γ, 100ng/mL TNF and 50ng/mL LPS. M2-type macrophage activation was achieved by supplementing growth media with 20ng/mL IL-13 and 20ng/mL IL-4. M1 macrophages display a rounded morphology, whilst M2 macrophages display a tortuous, dendritic morphology.

**Supplementary Figure 5:**
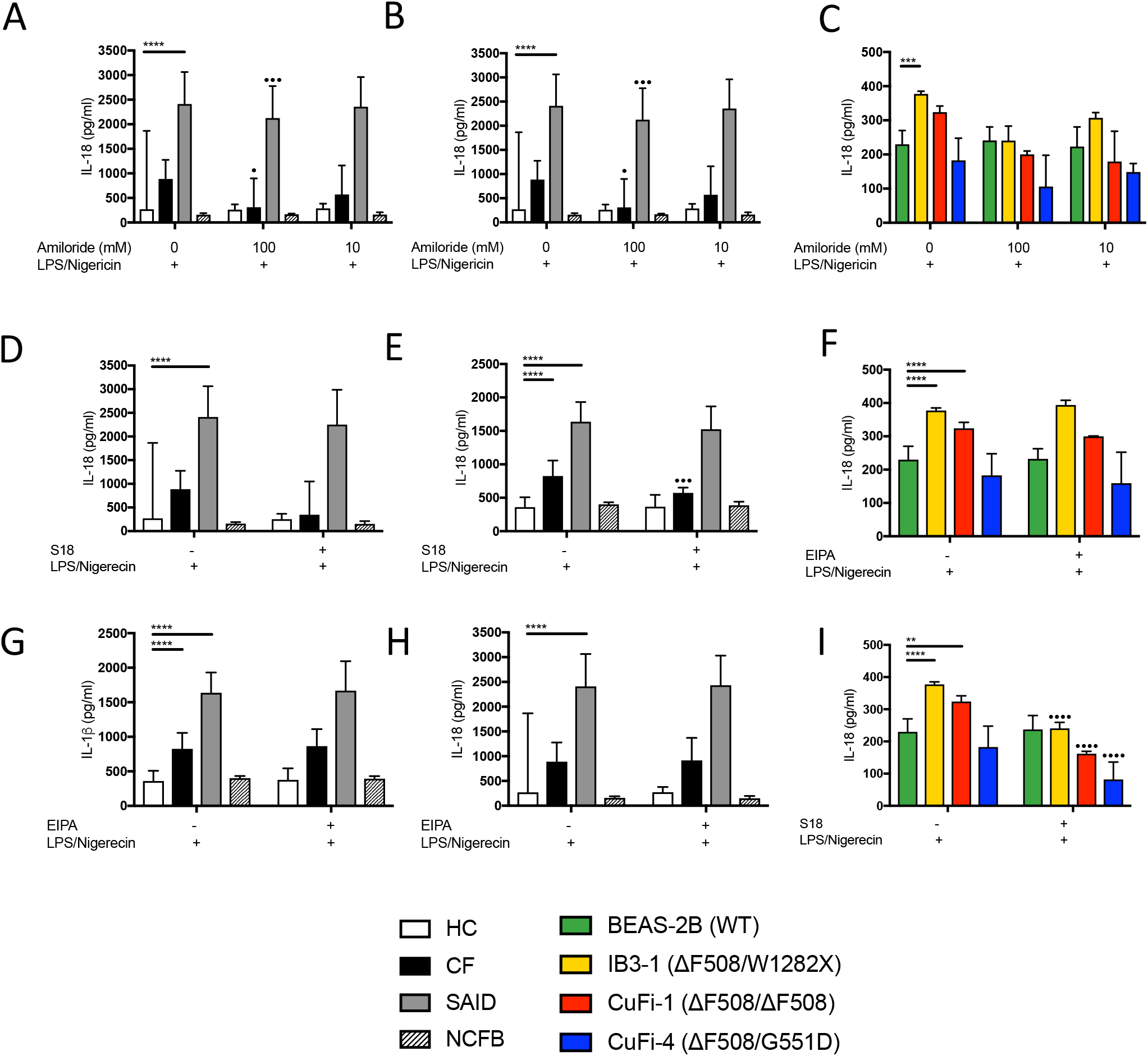
Inhibition of amiloride-sensitive sodium channels modulates inflammation in cells with CF-associated mutations. **A-I** ELISA assays were used to detect IL-18 (A, D, G) and IL-1β (B, E, H) in monocytes from HC (n=10), patients with CF (n=10), SAID (n=4) and NCFB (n=4) and IL-18 in HBECs (n=3) (C, F). Cell stimulation was as follows: Amiloride (100mM or 10mM, 1hour) (A-C), EIPA (10mM, 1 hour) (D-F), S18 derived peptide (25mM, 4 hours) (G-I) were used as a pre-treatment before a stimulation with LPS (10ng/mL, 4 hours) and Nigericin (1*μ*M) for the final 30 minutes. Data Information: A 2-way ANOVA statistical test was performed with Tukey post-hoc correction (p values * = 0.05, **= 0.01, ***= 0.001 and ****= 0.0001).

